# Mitochondrial DNA Density Homeostasis Accounts for the Threshold Effect in Human Mitochondrial Disease

**DOI:** 10.1101/078519

**Authors:** Juvid Aryaman, Iain G. Johnston, Nick S. Jones

**Affiliations:** Department of Mathematics, Imperial College London, London, United Kingdom; School of Biosciences, University of Birmingham, Birmingham, United Kingdom

## Abstract

Mitochondrial dysfunction is involved in a wide array of devastating diseases but the heterogeneity and complexity of these diseases’ symptoms challenges theoretical understanding of their causation. With the explosion of -omics data, we have the unprecedented ability to gain deep understanding of the biochemical mechanisms of mitochondrial dysfunction. However, there is also a need to make such datasets interpretable, and quantitative modelling allows us to translate such datasets into intuition and suggest rational biomedical treatments. Working towards this interdisciplinary goal, we use a recently published large-scale dataset, and develop a mathematical model of progressive increase in mutant load of the MELAS 3243A>G mtDNA mutation to develop a descriptive and predictive biophysical model. The experimentally observed behaviour is surprisingly rich, but we find that a simple, biophysically-motivated model intuitively accounts for this heterogeneity and yields a wealth of biological predictions. Our findings suggest that cells attempt to maintain wild-type mtDNA density through cell volume reduction, and thus energy demand reduction, until a minimum cell volume is reached. Thereafter, cells toggle from demand reduction to supply increase, upregulating energy production pathways. Our analysis provides further evidence for the physiological significance of mtDNA density, and emphasizes the need for performing single-cell volume measurements jointly with mtDNA quantification. We propose novel experiments to verify the hypotheses made here, to further develop our understanding of the threshold effect, and connect with rational choices for mtDNA disease therapies.

**Author Summary:** Mitochondria are organelles which produce the major energy currency of the cell: ATP. Mitochondrial dysfunction is associated with a multitude of devastating diseases, from Parkinson’s disease to cancer. Large volumes of data related to these diseases are being produced, but translation of these data into rational biomedical treatment is challenged by a lack of theoretical understanding. We develop a mathematical model of progressive increase of mutant load in mitochondrial DNA, for the mutation associated with MELAS (the most common mitochondrial disease), to address this. We predict that cells attempt to maintain the ratio of healthy mtDNA to cell volume by reducing their cell volume until they reach a minimum cell volume. As mutant load continues to increase, cells switch strategy by increasing their energy supply pathways. Our work accounts for large-scale experimental data and makes testable predictions about mitochondrial dysfunction. It also provides support for increasing mitochondrial content, as well as reduction in dependence upon mitochondrial metabolism via the ketogenic diet, as relevant treatments for mitochondrial disease.

## Introduction

Mitochondria are organelles known for their role in the production of ATP, the major energy currency of the cell. Their dysfunction is implicated in a host of diseases because of their role in biosynthesis [1] and energy supply, as well as their importance in cell death signalling [2], implicating them in diseases ranging from neurodegeneration [3] to cancer [4]. Fundamental understanding of these organelles and their dysfunction is therefore of far-reaching biomedical importance.

Mitochondria generate ATP by pumping electrons across their inner membrane, to generate an electro-chemical gradient, which is used by ATP synthase to convert ADP to ATP. The process of electron pumping is known as the electron transport chain (ETC), and this pathway of ATP generation is called oxidative phosphorylation (OXPHOS). Mitochondria also possess their own circular DNA (mtDNA), which are held in multiple copy number per cell. These genomes encode 13 proteins (which encode subunits of complexes I, III and IV of the ETC and ATP synthase), 22 tRNAs and 2 rRNAs. An important class of diseases which affect mitochondria are those which are caused by a mutation in mtDNA. The most common [5,6], and most studied, of these is MELAS (mitochondrial encephalomyopathy, lactic acidosis, and stroke-like episodes) syndrome, which is often associated with a mitochondrial tRNA mutation at position 3243A>G of the mitochondrial genome. Its incidence rate shows large regional variability, with prevalence of 1:6000 in Finland [7], to 1:424 in Australia [8]. tRNAs affected by the mutation cause amino-acid misincorporations during translation, generating defective mitochondrial protein, and defective respiration, when mutant load (or heteroplasmy) is high [9].

A common feature in many diseases associated with mutations of mitochondrial DNA, including MELAS, is the non-linear physiological response of cells and tissues to increasing levels of mtDNA heteroplasmy. In particular, cells appear to be able to withstand high levels of heteroplasmy without showing any significant metabolic or physiological defect. For instance, fibroblasts possessing the MELAS mutation were shown to have unaffected respiratory enzyme activity until mutant load exceeded around 60% [10]. Also, Chomyn *et al.* [11] showed that oxygen consumption of cells does not significantly reduce until MELAS heteroplasmy exceeds ~90%. This observation has been named the threshold effect (reviewed in [12]).

It has been argued that the threshold effect occurs because mitochondria possess spare capacity at the translational, enzymatic and biochemical levels, which are each able to absorb some degree of stress, and thus delay the phenotypic response of increasing heteroplasmy, until a particular threshold heteroplasmy is exceeded, which is typically large [12]. Within this picture, each physiological feature (such as enzymatic activity or oxygen consumption), may be expected to display step-like behaviour with respect to increasing heteroplasmy. Asynchrony of thresholds between different features, such as 60% for enzyme activity [10] and 90% for oxygen consumption [11], may be explained by spare capacity at intermediate levels: a biochemical threshold effect in this example, where metabolic fluxes are altered to compensate for fewer functional enzymes [12].

A recent study published by Picard *et al.* [13] established 143B TK^*−*^ osteosarcoma cell lines containing the MELAS 3243A>G mutation across the full dose response of mutant load. They measured a diverse array of features including RNA expression, protein expression, cell volume, growth rates, mitochondrial morphology and mtDNA content. The sheer diversity of data collected, across multiple levels of heteroplasmy, makes this an important dataset in understanding the threshold effect and mitochondrial dysfunction. Under the interpretation of the threshold effect presented above, one might expect a monotonic response to heteroplasmy, as spare capacity is depleted and the cell seeks alternative means of energy provision. However, Picard *et al*. observed complex multiphasic responses across numerous physiological readouts as heteroplasmy was increased [13].

The authors of that study identified four distinct transcriptional phases in the gene expression profiles of MELAS 3243A>G cells: 0%, 20-30%, 50-90% and 100% mutant load. They argue that continuous changes in heteroplasmy results in discrete changes in phenotype, because there exists a limited number of states that the nucleus can acquire in response to progressive changes in retrograde signalling [13]. In this work, by considering a distilled subset of the data from [13], and using simple, physically motivated arguments, we attempt to provide a simplified account of this dataset to gain better understanding of the consequences of this mutation and the threshold effect.

Our mathematical model suggests that cells attempt to maintain homeostasis in wild-type mtDNA density at low heteroplasmies, through reduction of cell volume and therefore cellular energy demand. We propose the existence of a single critical heteroplasmy, where cells are no longer able to maintain this homeostasis, and toggle from energy demand reduction to supply increase. In this regime, energy supply pathways are upregulated. Our model also identifies an additional bioenergetic transition, in excess of 90% mutant load, as cells become fully homoplasmic. We explore the possibility of reduced transcriptional activity in mutant mtDNAs/mitochondria, limited tRNA diffusivity, and a connection between cellular proliferation rate and cell volume, finding all of the above to have explanatory power. We propose new experiments to verify the novel hypotheses made here, to drive forward understanding of the threshold effect.

## Results

### Per-cell Interpretation of Omics Data Highlights Multiphasic Dynamics in Response to Heteroplasmic Load

We aim to understand mean cellular behaviour in response to rising levels of 3243A>G heteroplasmy. It is therefore important that the data we use to build this description is in per-cell dimensions. We perform normalisations of the data from [13] to create measures of overall gene expression of bioenergetic pathways from measurements of individual genes, and adjust for potential bias from variable cell volume induced by heteroplasmy (see Materials and Methods). Fig. 1 shows the core subset of physiological features from [13] we attempt to describe here, after these normalisations.

**Figure 1.**
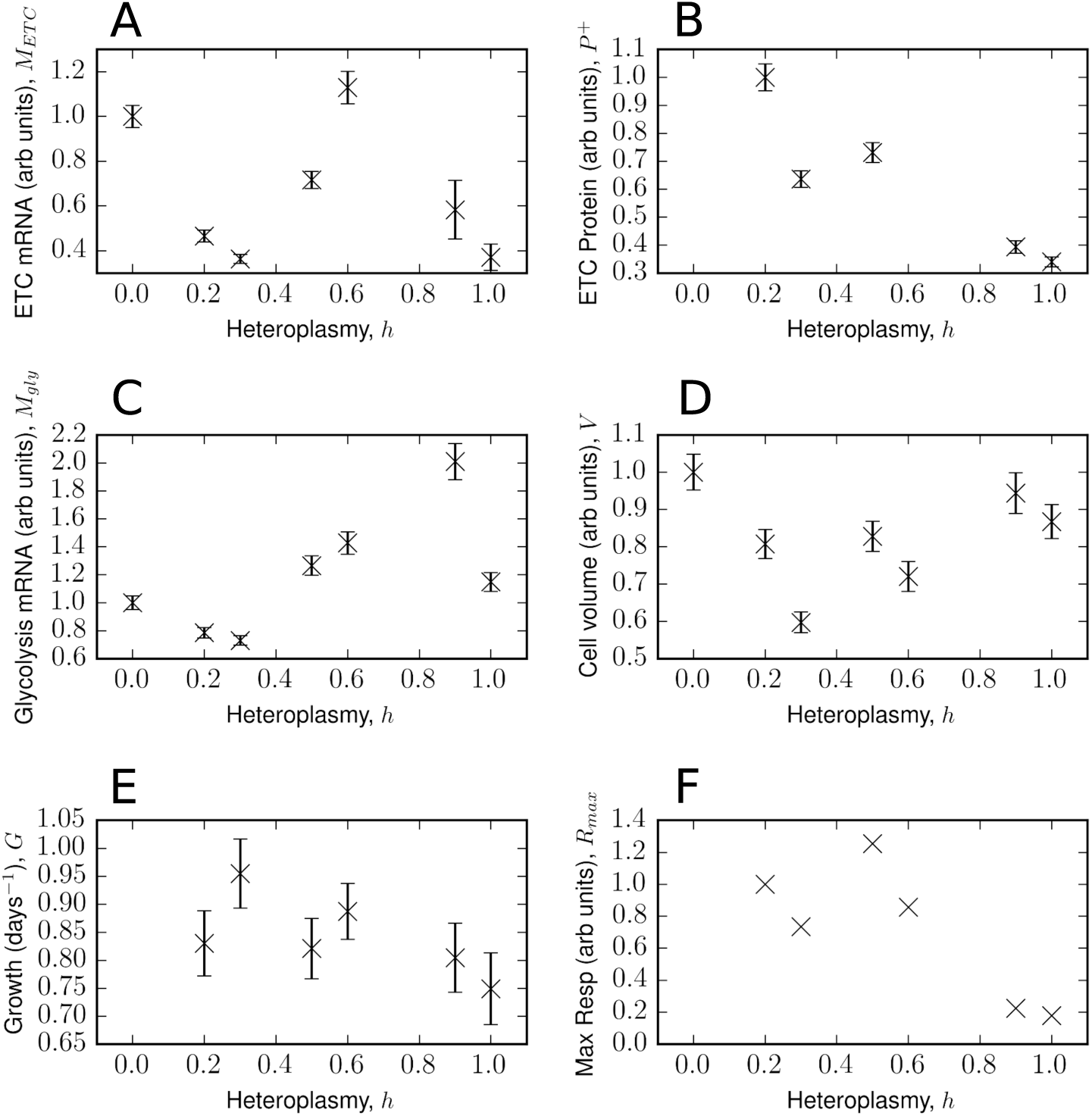
Multiphasic physiological response to increasing heteroplasmy – core data considered from Picard *et al*. renormalized to be in per-cell dimensions. Selected measurements of 143B TK^−^ osteosarcoma cells heteroplasmic in MELAS 3243A>G, from Ref. [13]. A. mRNA levels for 11 mitochondrially-encoded electron transport chain subunits (COX3, ND2, ND5, CYTB, ND3, ND6, COX1, ND4, COX2, ND1, ND4L). B. Protein levels for complexes I, III and IV. C. Glycolysis mRNA levels, for genes (PKM2, ENO1, PGAM4, PGK1, GAPDH, ALDOA, PFKP, GPI, HK2 and SLC2A1). All errors in A-C are the standard error of the transformed renormalised mean (see Data Normalization for details of how data for these genes are combined, and Error Propagation for associated errors). D. Mean cell volume of an asynchronous population of growing cells ± SEM. E. Growth rate, determined by linear regression (see Growth Rate Determination). Error is the standard error in the slope from linear regression F. Maximum respiratory capacity. See [13] for experimental protocols.

A simple interpretation of the threshold effect predicts the existence of spare capacity in the transcription, translation, enzyme complex and biochemical levels of the cell, in response to increasing heteroplasmy [12]. Under this interpretation of the threshold effect, we might expect all of these functions to have no more than one turning point with increasing heteroplasmy.

However, the data in Fig. 1, and indeed the dataset of Picard *et al*. [13] overall, shows a much more complex response. For instance, ETC transcripts clearly show two turning points, suggesting some kind of transient compensatory response. Across the features, these data also appear to be asynchronous in their turning points, for instance ETC transcripts peak at *h* = 0.6, but glycolysis transcripts peak at *h* = 0.9. This highlights the need for an extension in our understanding of the threshold effect, as well as the challenge in trying to parsimoniously model such a complex dataset.

We note that the measurement uncertainty, where reported in [13], for our selected features of interest are small relative to the variation with respect to heteroplasmy (see Fig. 1), justifying a non-linear fit to the data. It should be noted, however, that this uncertainty only reflects the technical variability in measurement, and does not include potential biological variability of these features. We use a Bayesian approach to appropriately account for this uncertainty, see Generative Model Description.

### Integrated Omics Data Motivates a Model of the Causal Relationships between Bioenergetic Variables

We present a qualitative description of our model in Fig. 2, which we will develop into a full quantitative description below. One of our central claims is the existence of a single transition in cellular behaviour, in response to increasing heteroplasmy of the 3243A>G mutation, over the 0-90% heteroplasmy range. We propose that, at low heteroplasmy values, cells attempt to maintain homeostasis in wild-type mtDNA density by reducing their volume. This reduces biosynthetic and translational energy demands, by the simplifying assumption that energy demand scales directly with cell volume.

**Figure 2.**
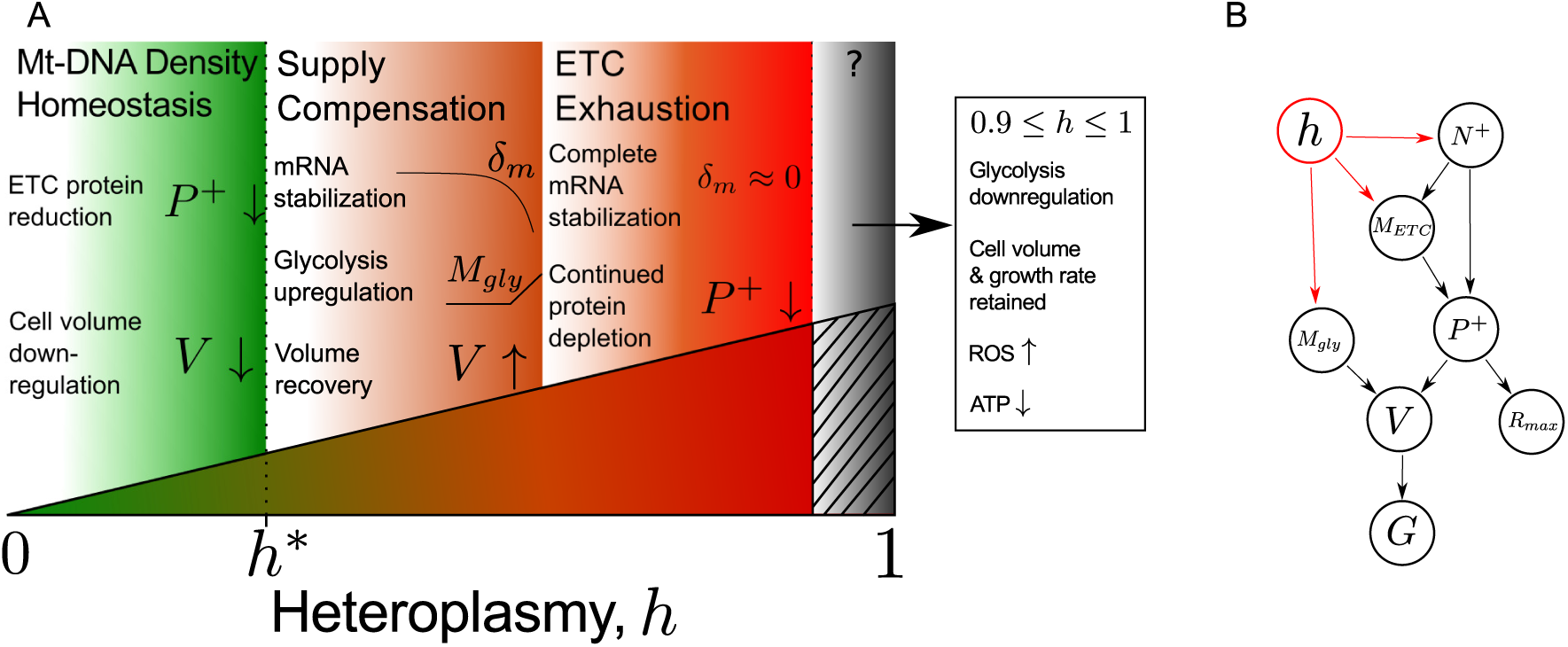
Qualitative description of continuous increase in MELAS mutant load. A. At low levels of heteroplasmy (*h*), cells attempt to maintain homeostasis in wild-type mtDNA density by reducing their volume (*V*). This reduces energy demands, allowing energy supply/demand balance to be maintained despite rising heteroplasmy. Further increase in heteroplasmy triggers a energy demand/supply toggle at a critical heteroplasmy *h*^*\**^, where energy provision pathways are upregulated. This includes upregulation of both oxidative phosphorylation, by reduction in mRNA degradation (*δ*_*m*_), and glycolysis transcripts (*M*_gly_). Cell volume consequently recovers. Further increase in heteroplasmy exhausts ETC stabilization, as *δ*_*m*_ → 0, and ETC protein (*P*^+^) continues to deplete. In the transition to homoplasmy, glycolysis and ATP levels reduce, and yet cell volume and growth rate are retained. In this regime, the mode of energy production is unclear. B. Flow of causality in our mathematical model. *N*^+^ = wild-type mtDNA copy number, *M*_ETC_=ETC mRNA, *R*_max_=maximum respiratory capacity, and *G*=cellular growth rate. Heteroplasmy shown in red connecting to variables with explicit dependence.

Our model suggests that at a critical heteroplasmy, *h*\*, cells undergo a demand/supply toggle where energy supply is upregulated. Electron transport chain (ETC) transcripts are stabilized through reduced degradation, and glycolysis is increased. This bioenergetic compensatory behaviour at intermediate heteroplasmies allows cell volume to recover.

As heteroplasmy continues to increase, we claim that degradation of ETC transcripts becomes negligible. Thus, further increases in heteroplasmy results in reduction in ETC protein content and ETC exhaustion ensues.

These behaviours are captured in the mathematical framework of our model. However, as cells transition from 90% to 100% mutant mtDNA, another transition in cellular behaviour appears to occur, according to the data of [13]. Cells downregulate glycolysis, and yet retain cell volume and growth rate. The mode of energy production in this case is unclear, and opens new questions as to the most relevant energy supplies and demands in homoplasmic cells (see Key Claims and Predictions of Biophysical Model of Heteroplasmy).

### Interactions between Bioenergetic Variables can be Cast as a Bottom-up Quantitative Model

We now present a quantitative description of our model, see Table 1, whose mechanistic interpretations will be more fully explored in Key Claims and Predictions of Biophysical Model of Heteroplasmy. Our model attempts to unify the experimentally-measured features of [13] within a simple, physically plausible, bottom-up cell biological representation. We stress that our choice of model structure was not developed independently of the data in [13]; hence, at the level of choice of model structure, we have limited control of over-fit. However, the uncertainty in its parameters, given the data and a set of priors (see Generative Model Description), was computed using Bayesian inference. So whilst our parametrization of the model has statistical control for uncertainty, we have not employed a statistical model selection framework. We believe this to be appropriate and practically unavoidable, as our objective is to yield a new reduced account for these heterogeneous data, present novel hypotheses and propose new experiments.

**Table 1.** Mathematical model of MELAS 3243 A>G mutation with progressive mutant load. See Table S2 for parameter descriptions. Heteroplasmy=*h*, mtDNA copy number=*N*.

| Description | Equation | Equation number |
| --- | --- | --- |
| Wild-type mtDNA, $N^+$ | $N^+ = N(1 - h)$ | (1) |
| ETC mRNA, $M_{\text{ETC}}$ | $M_{\text{ETC}} = \frac{\beta}{\delta_m + 1} N^+$ | (2) |
| | $\delta_m(h) = \frac{k_{\text{mRNA}}}{1 + \exp[k_m(h - h_0)]}$ | (3) |
| ETC protein, $P^+$ | $P^+ = \frac{N^+ M_{\text{ETC}}}{\delta_p}$ | (4) |
| Glycolysis mRNA, $M_{\text{gly}}$ | $M_{\text{gly}} = \begin{cases} c_1, & h \leq h^* \\ m_2 h + c_2, & h > h^* \end{cases}$ | (5) |
| Cell volume, $V$ | $\underbrace{k_o P^+ + k_g M_{\text{gly}}}_{\text{supply}} = \underbrace{V}_{\text{demand}}$ | (6) |
| Growth rate, $G$ | $G = \frac{k_{gr}}{V}$ | (7) |
| Maximum respiratory capacity, $R_{\text{max}}$ | $R_{\text{max}} = k_p P^+$ | (8) |

#### Wild-type mtDNA scaling

A theme apparent in the data of Picard *et al.* is an overall downward trend of ETC mRNA and ETC protein with increasing heteroplasmy (*h*). We therefore use the hypothesis that these quantities scale with the amount of wild-type mtDNA (*N*^+^). We assume that *N*=mtDNA copy number=const (set to 1 after normalisation, without loss of generality) which then defines *N*^+^, see Eq.(1). The successful performance of this simple model for *N*^+^ is shown in Figure S1. Note that this model has no free parameters, so was neglected in our Bayesian inference.

#### ETC mRNA

Transcript copy number is determined by the balance of transcription (*β*) and degradation (*δ*_*m*_) rates. Given our assumption of *N*^+^ scaling, it can be shown (see Text S1) that Eq.(2) may be used to model the ETC mRNA pool size (*M*_ETC_). We further assume a constant mean transcription rate *β* for parsimony, and allow the degradation rate *δ*_*m*_ to vary with heteroplasmy in response to cellular signals. We require the degradation rate to be high for low heteroplasmies, and low at high heteroplasmies, to describe the ability of cells to upregulate their transcript copy number with rising heteroplasmy. A biologically-motivated choice of function which achieves this is a sigmoid, see Eq.(3), where *k*_mRNA_, *k*_*m*_ and *h*_0_ are constants.

#### ETC protein

It is intuitive to assume that mean protein levels scale with transcript levels, although this relationship may be noisy [14]. Following a similar assumption for ETC mRNA, we also assume that ETC protein (*P* ^+^) scales with wild-type mtDNA levels. Using analogous arguments to *M*_ETC_ (see Text S1), we show that a reasonable model for ETC protein is Eq.(4), where *δ*_*p*_=const, denotes the baseline degradation of mitochondrial protein.

#### Glycolysis mRNA

We assume that the glycolysis mRNA pool size (*M*_*gly*_) is invariant to heteroplasmy, until a critical heteroplasmy *h*^*\**^, where glycolysis is gradually upregulated as a result of cellular control. It is therefore parsimonious to assume that glycolysis regulation obeys a spline of a constant and linear model which toggles at *h*^*\**^, see (5), where *c*_1_ and *m*_2_ are free parameters, and *c*_2_=*c*_1_ *− m*_2_*h*^*\**^, by continuity.

#### Cell volume

We propose that the energy demands of the cell may be well approximated as scaling with cell volume, see Text S1 for further discussion. As glycolysis and OXPHOS provide energy supply to first order, we assume that mean cell volume in an asynchronous population of cells (*V*) is effectively determined by a scaled sum of glycolysis and OXPHOS contributions to energy balance, such that the cell obeys an energy supply=energy demand relationship, see Eq.(6), where *k*_*o*_, *k*_*g*_=const.

From Fig. 1, it is clear that this assumption fails at *h*=1, where glycolysis levels and cell volume are comparable to *h*=0 levels, and yet ETC proteins are only 30% of wild-type levels. As ATP levels are below wild-type levels in these cells (see Fig. S6E in [13]), the mode of energy production is not clear and further metabolomic data may be required. We therefore exclude all *h*=1 data, and limit the domain of our model to 0 ≤ *h* ≤ 0.9.

#### Growth rate

We observe that the cellular proliferation rate (which we call growth rate, *G*, for consistency with [13]) varies with heteroplasmy (see Fig. 1). We hypothesize that there exists a relationship between mean cell volume and growth rate. It can be shown that, assuming individual cells increase their volume linearly through the cell cycle, growth rate varies inversely with mean cell volume (shown in Text S1). This is shown in Eq.(7), where *k*_*gr*_ is a constant.

We show in Text S1 that, under an exactly exponential model of cell growth, *G* is independent of *V*. However, given that there is presumably a wide class of cytoplasmic growth-vs-time profiles which cells may obey, we use a linear model as a parsimonious example of how cell growth may be connected to cytoplasmic volume.

#### Maximum respiratory capacity

It has long been recognised that cells carrying the MELAS mutation experience a respiratory defect when heteroplasmic load exceeds approximately 90% [11, 15], and that this is due to a defect in protein synthesis [9, 11]. We therefore assume that maximum respiratory capacity (*R*_max_) is always determined by protein content. This yields a simple linear expression, see Eq.(8), where *k*_*p*_=const.

#### Model summary

In summary, our model of mean cellular behaviour with respect to heteroplasmy describes 7 features from Picard *et al.* [13] (*N*^+^*, M*_ETC_*, P* ^+^*, M*_gly_*, V, G, R*_max_) and has 12 adjustable parameters (as discussed later this is fewer than the number required for 7 linear models), a table of which is shown in Table S2. In writing down this phenomenological model, we have attempted to account for a physiologically important subset of the data generated in [13], using bottom-up arguments wherever possible. In doing so, a number of novel, falsifiable, hypotheses are made.

## Parametrizations of a Simple Biophysical Model Account for Complex Observations Across Range of Heteroplasmic Load

The fit of the model described above is shown in Fig. 3. Between 0 ≤ *h* ≤ *h*^*\**^, *h*^*\**^ being the critical heteroplasmy where glycolysis is upregulated (0.34 ≤ *h*^*\**^ ≤ 0.44, 25-75% CI), our model reproduces the reduction in ETC transcript pool size. Similarly, we observe that ETC protein pool size also reduces, as does cell volume and maximum respiratory capacity.

**Figure 3.**
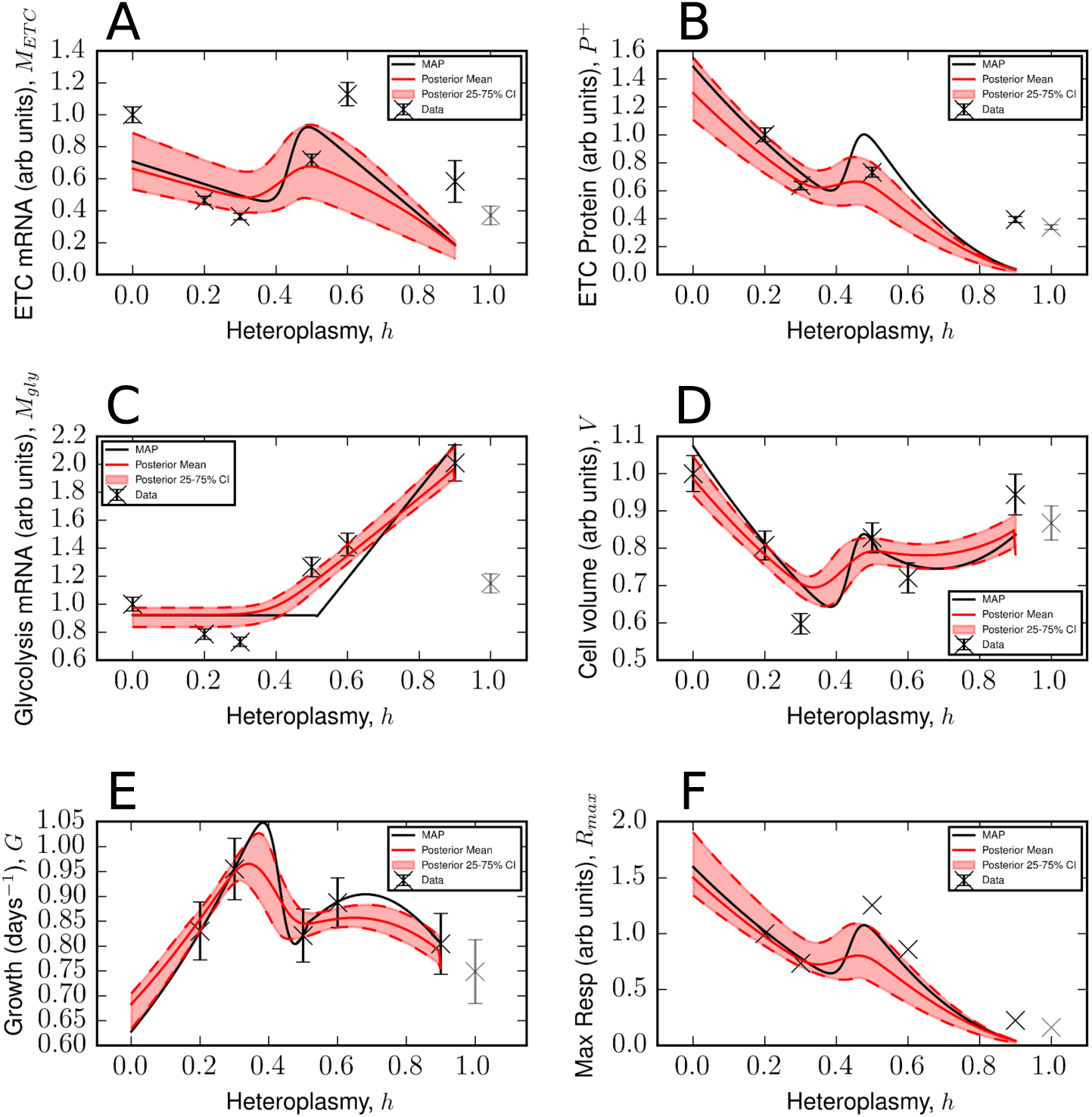
A simple biophysical model is consistent with complex observations across range of heteroplasmic load A-F. Approximations for the maximum a posteriori estimate (black line), posterior mean (red line) and 25-75% confidence intervals (pink bands), for the model fits to selected data from [13]. The model makes predictions over the range 0 ≤ *h* ≤ 0.9, see Main Text. Data for *h*=1 have been plotted in grey, as they have not been used to train the model. Error bars are conservative and merely show the technical variability reported in [13], see Materials and Methods.

Our model is able to successfully capture the transient compensatory responses in ETC mRNA, ETC protein and cell volume which begin around the critical heteroplasmy *h*^*\**^. For heteroplasmies between 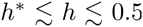, ETC mRNA degradation reduces causing ETC mRNA to be upregulated, along with ETC protein and maximum respiratory capacity. In this region, glycolysis becomes induced above wild-type levels, and cell volume can be observed to also recover.

In excess of *h* ≈ 0.5, our model shows the observed reductions in ETC mRNA, ETC protein and maximum respiratory capacity. We see that continued upregulation of glycolysis mRNA allows cell volume to remain at an approximately constant value, although diminished relative to a wild-type cell. Consequently, heteroplasmic cells between 0.2 ≤ *h* ≤ 0.9 are predicted to proliferate at a faster rate than wild-type cells (see Fig. 3E).

## Key Claims and Predictions of Biophysical Model of Heteroplasmy

Here, we revisit the interpretations of our model in light of the mathematical description developed above, and explore the evidence for the biological insights it provides. We make experimental proposals to validate our claims, which are given in Text S2. The set of mechanistic interpretations which follow from our mathematical model, see Fig. 4, are:

- Wild-type mtDNA density is maintained homeostatically at low heteroplasmy
- There exists a minimum cell volume which is approached at the critical heteroplasmy
- Cells toggle from demand reduction (i.e. cell volume reduction) to supply increase (i.e. glycolysis and ETC mRNA upregulation), at the critical heteroplasmy
- Mutant mtDNAs do not significantly contribute to the mitochondrial mRNA pool
- Mitochondrial tRNAs remain moderately localised to their parent mtDNA
- Maximum respiratory capacity is determined by ETC protein levels through a linear relationship
- Cell growth rate is the reciprocal of mean volume, thus smaller cells grow faster

**Figure 4.**
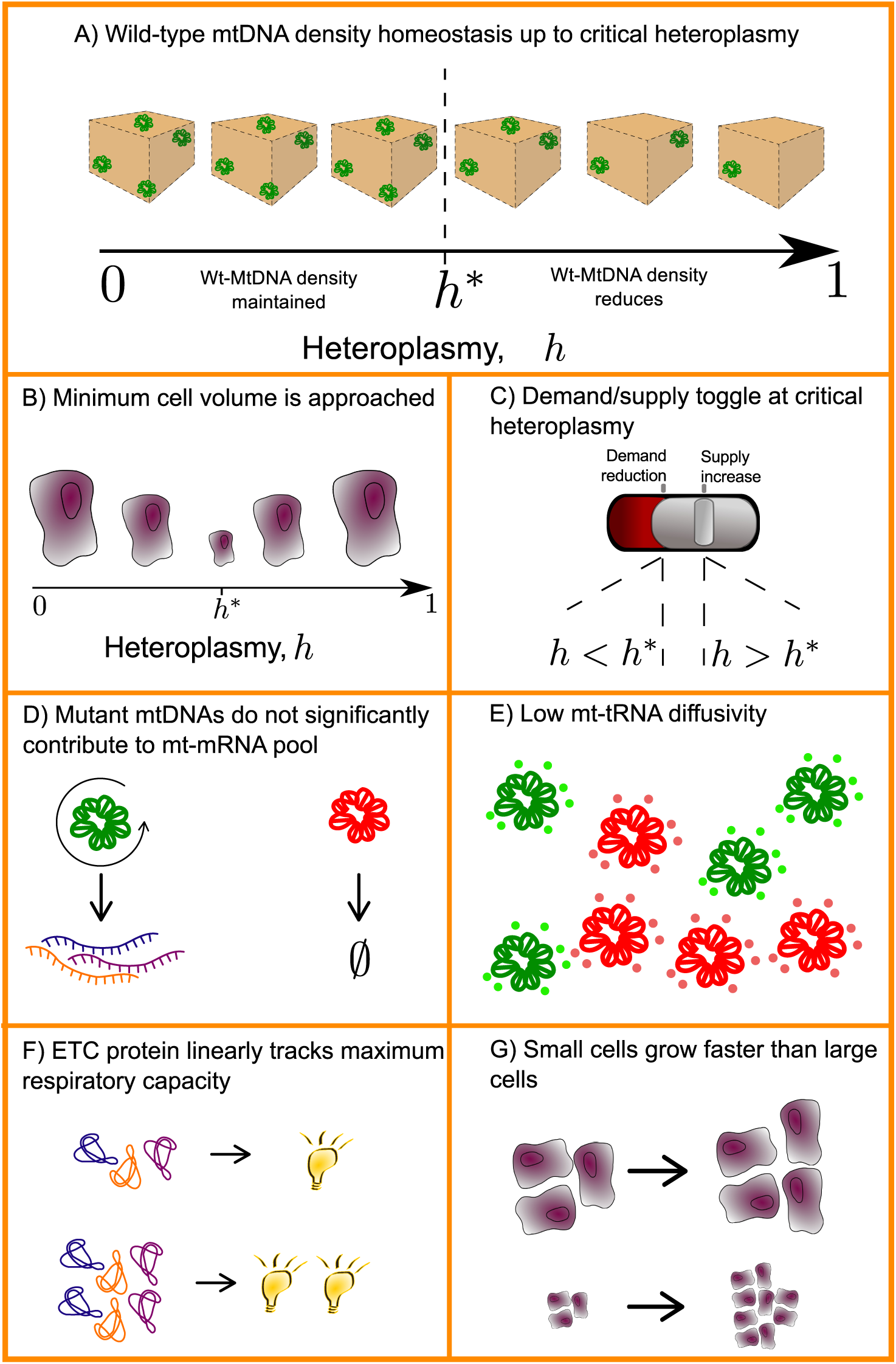
Claims and interpretations of biophysical model. A. Wild-type mtDNA density is maintained by the cell when heteroplasmy is below the critical heteroplasmy, *h*^*\**^. B. Cells achieve this through reduction of their cell volume, and therefore energy demand, as wild-type mtDNA copy number diminishes. This continues until a minimum cell volume is reached at *h*^*\**^. C. At heteroplasmies above *h*^*\**^, wild-type mtDNA homeostasis cannot be maintained through volume reduction due to the existence of a minimum cell volume. Cells therefore switch their strategy to increase energy supply. D. Mutant mtDNAs have a much lower contribution to the transcript pool than wild-type mtDNAs. E. tRNAs stay local to their parent mtDNA, meaning that mRNA must come into contact with a wild-type mtDNA to be translated into normal protein. F. Maximum respiratory capacity is linearly proportional to ETC protein. G. Assuming that cytoplasm grows linearly through the cell cycle, smaller cells proliferate faster than larger cells.

### Wild-type mtDNA density homeostasis is maintained until a minimum volume is reached near the critical heteroplasmy

The parameter *h*^*\**^ determines the extent of mutant load, for which the cell begins to upregulate ETC mRNA and glycolysis mRNA. But what causes this change in behaviour, at this particular value of heteroplasmy? By examining the posteriors of our model fit (Fig. 3) we infer that cell volume takes its minimum value shortly before the most probable value of *h*^*\**^ (see Fig. 5). We hypothesize that an attempt to conserve wild-type mtDNA density (*N* ^+^/*V*) determines the position of *h*^*\**^.

For 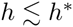, wild-type mtDNA density is maintained despite increasing heteroplasmy, because cell volume diminishes. As a result of this reduced demand, the cell can tolerate diminished mitochondrial power supply. However, cell shrinkage cannot continue indefinitely and we hypothesize that the cell reaches a minimum cell volume at *h* ≈ *h*^*\**^. Once heteroplasmy exceeds this value, the cell toggles its energy balance strategy from demand reduction to supply increase, and the cell recovers in volume.

**Figure 5.**
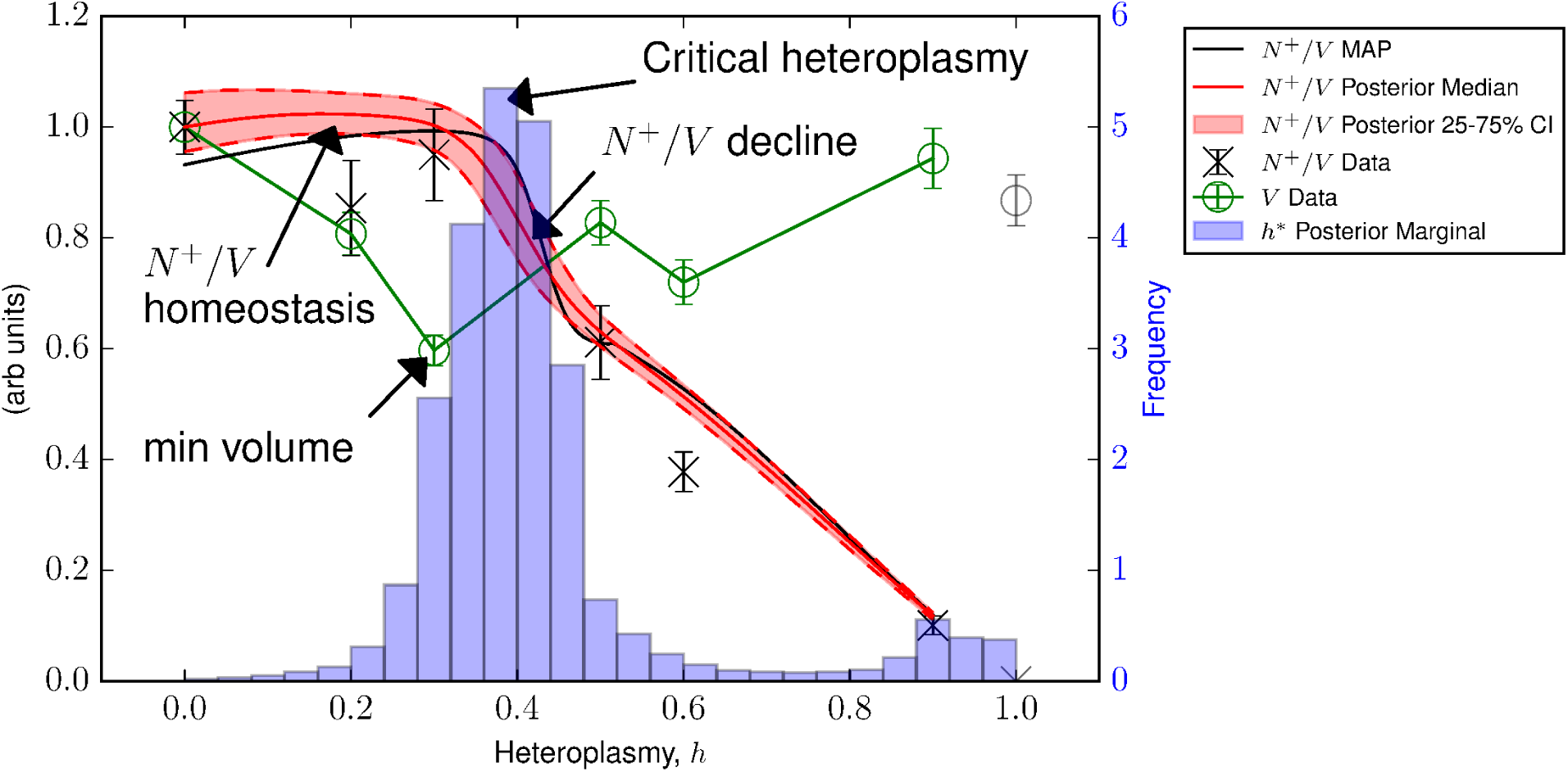
Wild-type mtDNA density (*N* ^+^/*V*) homeostasis may trigger supply/demand toggle. Posterior statistics show an initial maintenance of *N* ^+^/*V*. When cell volume (*V*) takes its minimum value, the most probable value of *h*^*\**^ shortly follows. *N* ^+^/*V* then reduces. We suggest that the inability of the cell to maintain *N* ^+^/*V*=const, due to the existence of a minimum cell volume, causes cells to toggle in their strategy at *h*^*\**^, from demand reduction to supply increase. Data from [13].

There is evidence in the literature that wild-type mtDNA density is an important quantity. Bentlage and Attardi [15] observed that long-term culture of heteroplasmic MELAS cells resulted in an increase in mtDNA copy number, resulting in increased oxygen consumption. Whilst this was often accompanied by a decrease in heteroplasmy, some cell lines also exhibited this at constant heteroplasmy. This is consistent with the cell attempting to increase the absolute number of wild-type mtDNAs, perhaps to compensate for heteroplasmic load, and suggests that the absolute value of *N* ^+^ is a physiologically important quantity.

The density of mitochondrial content per unit cytoplasmic volume has been observed by many authors to be tightly regulated and physiologically predictive. The historical observations of Posakony *et al.* [16] showed that the mean ratio of mitochondrial content to cytoplasmic volume is kept relatively constant throughout the cell cycle in HeLa cells, occupying ~10-11% of cytoplasmic area throughout. Similar observations have been reproduced in more recent studies, in various other systems. Rafelski *et al.* found in budding yeast that mitochondrial content was proportional to bud size, and that all buds attain the same average ratio regardless of the mother’s age or mitochondrial content [17], suggesting a stable scaling relation. Also, Johnston *et al.* [18] found that the density of mitochondrial mass was predictive of cell cycle dynamics, indicating that *N/V* (*N*=total number of mtDNAs) is physiologically relevant and potentially linked to cell energy supply and growth dynamics. Indeed Jajoo, Paulsson and co-workers [19] found that the density of mitochondrial DNA tracks the quantity of cytoplasm inherited upon division in wild-type fission yeast. Finally, Otten *et al*. found a positive correlation between cell volume and mtDNA copy number in zebrafish oocytes [20].

We may speculate as to the interpretation of a minimum cell volume. One straightforward interpretation is that a minimum cell volume corresponds to a mechanical constraint: a cell may only become so small because the machinery required to perform tissue-specific metabolic and structural tasks require a minimum amount of space.

An alternative to this is a bioenergetic minimum cell volume. Numerous historical studies have shown that there exist appreciable energy demands which do not scale linearly with volume [21–23]; for instance, processes which only serve the nucleus such as DNA-replication, or demands associated with the plasma membrane. If a unit volume of cytoplasm has a particular energy output, which satisfies the energy demand of that unit volume plus an energy surplus, then continued reduction of cell volume results in the total energy surplus of the cytoplasm being unable to meet the demands of the nucleus and plasma membrane. At this bioenergetic minimum cell volume, the nucleus may signal to increase energy production pathways to restore the energy surplus of a unit volume of cytoplasm. If we assume that power supply per unit volume must be maintained, then as cells become smaller in radius (*r*), surface area demands per unit volume scale with *r*^2^/*r*^3^=1/*r* whereas constant energy demands per cell per unit volume scale with 1/*r*^3^. In this way, demands associated with cell surface area may be the first to become prohibitive as cells reduce in size, more so than constant demands which scale with a larger negative power of *r*. Both mechanical and bioenergetic limits no doubt exist, but which of these constraints is first encountered upon volume reduction is open.

### ETC mRNA degradation diminishes at the critical heteroplasmy contributing to energy demand/supply toggle

The induction of glycolysis at the critical heteroplasmy is observed in our model by construction, see Eq.(5), since glycolysis is modelled to increase linearly when heteroplasmy exceeds this point. However, by observing the posterior distribution of the ETC mRNA degradation rate (see Fig. 6), we see that the critical heteroplasmy also coincides with the beginning of reduction in ETC transcript degradation with respect to heteroplasmy. Since ETC mRNA pool size varies with the inverse of this degradation rate (see Eq.(2)), ETC transcripts are consequently upregulated in tandem with glycolysis transcripts. This occurs until ETC degradation diminishes to negligible levels around *h* ≈ 0.5, where this particular control mechanism becomes exhausted. Thus, the critical heteroplasmy coincides with a shift from energy demand reduction, to supply increase from both glycolysis and OXPHOS contributions.

**Figure 6.**
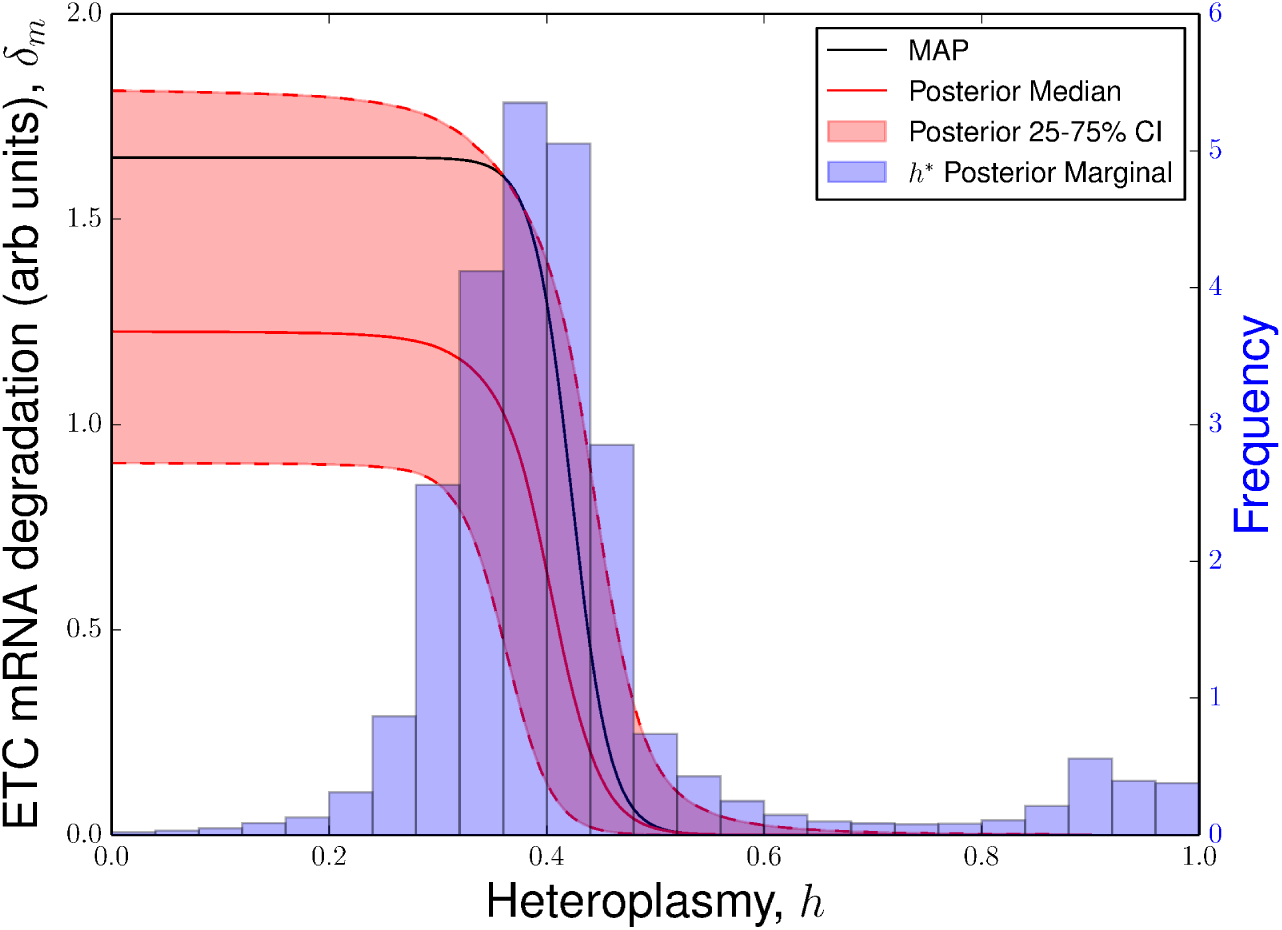
Critical heteroplasmy induces ETC mRNA stabilization. Posterior distributions for ETC mRNA degradation rate and critical heteroplasmy (0.34 ≤ *h^*^ ≤* 0.44, 25-75% CI and 0.27 ≤ *h*^*\**^ ≤ 0.89 5-95% CI). It can be seen that the critical heteroplasmy coincides with the reduction in ETC degradation, signalling an energy demand/supply toggle.

Since mtDNA is transcribed as a single polycistronic transcript [24], the stoichiometry of individual mRNA species must be controlled via active degradation. This is achieved by a balance between processes which stabilize and degrade mRNA [25]. The Picard data set can be explored further to seek corroborating evidence, by observing the ratio of ETC mRNA degraders to stabilizers. We find a qualitative similarity between this ratio (see Figure S7) and the posterior distribution of the ETC degradation rate (see Fig. 6), both displaying a substantial reduction between *h*=0.3 and *h*=0.5.

### Mutant mtDNAs do not significantly contribute to the mitochondrial mRNA pool

Our hypothesis that ETC mRNA transcript pool size is proportional to wild-type mtDNA copy number, i.e. *M*_ETC_ ∝ *N* ^+^, was invoked as a simple explanation for the overall downward trend with heteroplasmy. We favoured this explanation over, for instance, allowing the transcript birth rate to decrease with heteroplasmy, as such behaviour would contradict the behaviour of the degradation rate which acts to increase transcript pool size. The implication of this model is that mutant mtDNAs do not contribute strongly to the transcript pool, either through a transcription defect or selective degradation of all transcripts from mutated mtDNAs, the precise mechanism is not prescribed by the model. (We discuss alternative models to *N* ^+^ scaling for *M*_ETC_ in Alternative Hypotheses)

### Mitochondrial tRNAs are relatively localised to their parent mtDNA

It has been observed that homoplasmic MELAS cells are able to translate mitochondrially-encoded proteins; however, misincorporations cause these translation products to become unstable [9]. Assuming rapid degradation of such proteins, mutant mtDNAs are not expected to contribute strongly to the total ETC protein content of the cell, which has been confirmed by experimental observation in homoplasmic cells [9]. We adopted the model of ETC protein being proportional to wild-type mtDNA copy number, i.e. *P* ^+^ ∝ *N* ^+^, as a parsimonious model for such scaling.

One interpretation of this assumption is that we can identify Eq.(4) as obeying mass-action kinetics between ETC mRNA and wild-type mtDNA molecules, with a constant baseline degradation rate. A simple interpretation of this, is that ETC mRNAs must come into the proximity of wild-type mtDNAs, to be translated.

One way in which this might be achieved is if tRNAs remain spatially localised to their parent mtDNA; in other words, tRNAs have low diffusivity. ETC mRNAs which come into contact with mutant mtDNAs are translated into mutated protein only, since mutated tRNAs are much more available, which are then rapidly degraded. Conversely, mRNAs which localise with wild-type mtDNAs are only translated into normal protein, since only wild-type tRNAs are available. (We discuss alternative models to *N* ^+^ scaling for *P* ^+^ in Alternative Hypotheses)

Evidence in the literature for this claim is mixed. It has been observed that mitochondrial mRNAs, such as ND6, localise to mtDNA suggesting that mtDNA may be a site for mitochondrial translation [26]. However, cybrid experiments involving homoplasmic tRNA mutants 3243A>G and 4269A>G are able to recover their respiratory function by fusing such cells together to form hybrids [27]. Their recovery is presumably due to the diffusion of the healthy form of each tRNA, so that normal proteins may be translated. See Text S2 and Table S1 for experimental suggestions to determine the extent of tRNA diffusivity.

### Cell volume is not explained by cell cycle variations

Our model predicts that cells, on average, change their size as heteroplasmy is varied, due to variation in power supply from OXPHOS and glycolysis. However, since cells vary their volume by a factor of 2 throughout the cell cycle, it is possible that cells with different heteroplasmies spend different durations at various stages of the cell cycle, explaining the observed variation in expected cell volume with heteroplasmy (see Fig. 1D). We sought evidence for this hypothesis, by computing the ratio of the expression level, for genes associated different stages of the cell cycle [28], see Figure S8. However, we found little evidence to support the enrichment of cell cycle markers at any particular level of heteroplasmy.

### OXPHOS contributions to energy supply are stabilized at the critical heteroplasmy

The relative contribution of OXPHOS to energy supply, i.e. *k_o_P* ^+^/(*k_o_P* ^+^ + *k*_*g*_*M*_*gly*_), is also interesting to observe as heteroplasmy is varied. We observed a transient stabilization in OXPHOS contributions around *h*^*\**^. A discussion of this is presented in Text S3.

### Cells proliferate inversely with their size

Due to our reciprocal model connecting cell volume and growth (see Eq.(7)), our model suggests that wild-type cells proliferate more slowly relative to heteroplasmic cells due to their larger size.

### Maximum respiratory capacity linearly tracks ETC protein content

It has long been suggested that cells above a particular threshold heteroplasmy experience a respiratory defect [10, 11, 15]. In our model, we found that a simple linear relationship between ETC protein and maximum respiratory capacity was sufficient to describe the data available (see Eq.(8)). With a more classical interpretation, we might have expected the need to deploy a model which has switching behaviour in excess of 60% heteroplasmy [10] for maximum respiratory capacity, in analogy with glycolysis transcript levels (see Eq.(5)).

### Reactive oxygen species may explain the transition to homoplasmy but mode of energy production remains unclear

In Eq.(6), we claim that cell volume is determined by the weighted sum of glycolysis transcripts and ETC protein. Over the range 0.9 < *h* ≤ 1, glycolysis transcripts reduce by 57%, whereas ETC protein and cell volume remain comparable, thus breaking the supply=demand relationship, as we have modelled it. Consequently, our model fails to describe the transition from *h*=0.9 → 1.

A potential explanation for the reduction in glycolysis transcripts over this range, comes from the fact that glycolysis provides substrate for oxidative phosphorylation. Damaged electron transport chain proteins may produce an excess of reactive oxygen species (ROS) [29], which can damage mitochondrial proteins, DNA and membranes. If, at high heteroplasmy, any flux through the electron transport chain causes high levels of ROS, then cells may attempt to reduce flux through glycolysis, to avoid production of these species.

Some evidence from Picard *et al.* supports this hypothesis, where superoxide dismutase (SOD) activity is largely constant with heteroplasmy, except for homoplasmic mutant MELAS cells, which have ~ 20% higher SOD activity than wild-type cells (see Fig. S7D of [13]). Furthermore, it is known that ROS can reversibly inhibit the activity of GAPDH, one of the enzymes involved in glycolysis [30, 31].

However, given that fatty acid oxidation (see Figure S9) is strongly downregulated over this range, it remains unexplained how homoplasmic mutant cells maintain their cell volume (and growth rate), given their reduced reliance upon mitochondrial and glycolytic metabolism. Further metabolomic measurements may be required to uncover this mode of energy production.

ATP levels are also observed to decrease over this range (see Fig. S6E of [13]), which may even suggest that an alternative fuel currency besides ATP supports the growth and size of these cells. However, more careful investigation of this observation may be of value, since it is important to draw the distinction between ATP pool sizes and ATP fluxes, the latter perhaps being more indicative of ATP usage, and the former being indicative of only relative production/consumption rates.

## Alternative Hypotheses

Here we explore several alternative hypotheses, and point out a number of reservations in accepting these alternatives over the model presented above.

### Wild-type mtDNA copy number scaling for ETC mRNA

Our claim that ETC mRNA scales with wild-type mtDNA copy number linearly (see Eq.(2)) was used for parsimony, but other models exist. We know that mtDNA exists in multiple copy number per organelle, one estimate is that there exists between 4-40 mtDNAs per mitochondrion [32]. If we use the model that mitochondria containing only mutated mtDNA are unable to transcribe, and mitochondria are otherwise able to transcribe, then this results in a non-linear scaling of *M*_ETC_ with *h*. Instead of scaling with *N* ^+^, ETC mRNA would scale with the probability that 100% of mtDNAs per organelle are mutated (*p*_100_, which follows a binomial distribution with probability *h*). We might think of this as an organellar threshold effect. This model is plausible if we assume that mitochondria power their own transcription, and even a single mtDNA is able to power transcription for the entire organelle.

However, since there exists a range of possible values for the number of mtDNAs per mitochondrion in a cell, this non-linearity is likely to be somewhat smoothed out. So having ETC mRNA scaling with *N* ^+^ is still plausible at a cellular level, even under this hypothesis, and is more parsimonious.

We also investigated the possibility that ETC mRNA scales with *N* ^+^ + *μN* ^*−*^, where *N* ^*−*^=the number of mutant mtDNAs (see Text S4), where we constrained 0 ≤ *μ* ≤ 1. We found large support for values of *μ* close to 1 (see Figure S3), but draws from the posterior distribution of *M*_ETC_ were often purely linear and thus inappropriate for understanding threshold effects (see Figure S15). For this reason, we rejected the mutant transcription model in favour of the model presented in the main text.

### Wild-type mtDNA copy number scaling for ETC protein

A similar organellar threshold argument of scaling with the probability that 100% of mtDNAs per organelle are mutated, *p*_100_ scaling (instead of *N* ^+^), may hold if mitochondria power their own translation. This would relax the need to invoke low tRNA diffusivity. But, again, for parsimony we favoured *N* ^+^ scaling.

We also explored the ability of misincorporation effects to explain the ETC protein data, as opposed to tRNA localisation (see Text S4). If tRNAs are well mixed, then an ETC protein may possess a tolerance to the number of misincorporations per protein. Upon performing Bayesian inference, we found that the most likely tolerance to the MELAS mutation was 100% of residues: in other words, ETC proteins are immune to the MELAS mutation. This has been shown experimentally to be incorrect [9]. Furthermore, draws from the posterior distribution of *M*_ETC_ were often purely linear and thus inappropriate for understanding threshold effects (see Figure S15). For these reasons, we rejected the tRNA misincorporation model in favour of the model presented in the main text.

## Discussion

In this study, through use of a distilled subset of data from [13] and using minimal arguments, we have attempted to explore the apparent marked difference between the complex multiphasic observations of Picard *et al*., and the classical step-like models associated with the threshold effect.

We argue that a single critical heteroplasmy, *h*^*\**^, is sufficient to explain this subset of data over the heteroplasmy range 0 ≤ *h* ≤ 0.9 and that other multiphasic behaviour arises naturally from the simple physical/biological assumptions of our model. Our model suggests that cells undergo an energy demand/supply toggle at *h*^*\**^, from demand reduction to supply increase. We hypothesize that homeostasis in wild-type mtDNA density is maintained via cell volume reduction, ensuring that the available functioning power sources are matched to a corresponding level of cellular demand, until a minimum cell volume is reached which coincides with *h*^*\**^. This triggers the demand/supply bioenergetic toggle where energy production pathways are upregulated. We believe this re-emphasizes the need for quantification of single-cell mtDNA content to be associated with volume measurements of the same cell: mtDNA density is a relevant physiological variable [16–20]. We find that the mode of energy production over the range 0.9 ≤ *h* ≤ 1 is unclear, and that further metabolomic investigations may be required to determine this.

Our model further generates hypotheses that mutant mtDNAs (or alternatively homoplasmic organelles) have a reduced contribution to transcription, that tRNAs have low diffusivity, and that a relationship exists between mean cell volume and cell growth. We have proposed novel experiments to verify these hypotheses, to further develop our understanding of the threshold effect.

A potential consequence of our predictions is that controlling either mtDNA copy number, wild-type mtDNA copy number, or wild-type mtDNA copy number density, to ensure optimal values of wild-type mtDNA copy number density could be valuable control axes in therapy. Increasing mitochondrial DNA copy number, for instance through activation of the PGC-1*α* pathway, may facilitate the increase of cell volume, deferring the critical heteroplasmy to higher values by delaying the approach towards a minimum cell volume. We might reason that this enhances a wild-type phenotype at higher heteroplasmy values, potentially deferring the full MELAS phenotype to higher heteroplasmies, which typically appears between ~ 50-90% mutant load [33]. Indeed, it has been found that increasing mitochondrial biogenesis can ameliorate mitochondrial myopathy in vivo [34].

We might also argue that as cells toggle from energy demand reduction to supply increase, further bolstering of this compensatory response may have clinical significance. For instance, since we observe that cells switch to glycolytic metabolism to compensate for diminishing mitochondrial power supply, further encouragement of this energy mode may be therapeutic. This is supported by the recent observation that promoting the hypoxia response is protective against multiple forms of respiratory chain inhibition [35]. Alternatively, since we predict that cells innately downregulate ETC mRNA degradation, seeking to upregulate mitochondrial transcription may aid the cell in maintaining a sufficient mRNA pool size. Furthermore, promoting alternative energy production pathways such as fatty acid oxidation via the ketogenic diet may also aid in reducing the dependence on oxidative phosphorylation. This diet has been associated with increased mitochondrial transcripts [36], mitochondrial content [36,37] and has been shown to slow mitochondrial myopathy progression in transgenic Deletor mice [37]. Indeed, the diet has recently been used in clinic as an adjunctive therapy for a patient suffering from MELAS, harbouring the 3260A>G mutation, which successfully decreased the frequency of seizures and stroke-like episodes [38].

## Materials and Methods

### Data Normalization

Bioenergetic pathways, such as glycolysis or oxidative phosphorylation, consist of a set of enzymes, whose corresponding genes may be correlated in their expression. To have some measure of the overall expression level of a pathway 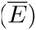, we use the mRNA concentration (in RPMK, reads per kilobase of transcript per million mapped reads), for each gene corresponding to enzymes of the pathway (*e*_*i,k*_(*h*), for gene *i* and technical replicate *k* at heteroplasmy *h*) and take a normalized sum

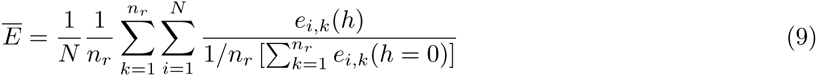

where *n*_*r*_=number of technical replicates, and *N*=number of genes in the pathway of interest. This quantity normalizes the expression level of each gene to *h*=0 levels, to avoid effects from consistently highly-expressed genes. The factor of 1/*N* results in 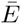 having the value of 1 at *h*=0, so may be interpreted as a fold-change in expression relative to *h*=0.

The standard error of 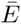 is given by

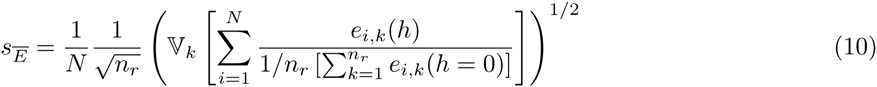

where 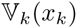 is the sample variance over *x*_*k*_. Eq.(9) and Eq.(10) are applied to glycolysis and ETC mRNAs in our main model, which yield dimensionless, normalized, measures of transcript levels for each biological pathway.

ATP synthase is excluded from both mRNA and protein data, as it is expected to be regulated differently from other ETC proteins. This difference arises because mitochondrial membrane potential is required for cell growth [39], and glycolytic ATP may be used, even in cells without mtDNA, by ATP synthase to maintain membrane potential [40]. Thus, protein levels of ATP synthase may be expected to be regulated quite differently to those of the electron transport chain, and not generally indicative of respiratory activity.

For ETC protein, we simply use the sample mean of complexes I, III and IV, since the data given by Picard *et al.* [13] is already normalized.

### Data Transformation to Per-Cell Dimensions

The data we consider of Picard *et al.* [13], consists of RNA-seq and Western blot measurements for mRNA and protein levels respectively. We wish to model the bioenergetic strategy of an average cell, so it is important that the data we use to parametrize our model is of per-cell dimensions. We show in Text S5 that it is appropriate to multiply protein and transcript data by cell volume to gain per-cell dimensions.

### Error Propagation

Our work focuses on describing mean behaviour with respect to heteroplasmy, so uncertainty in this mean must be quantified. For *M*_gly_ and *M*_ETC_, we used error propagation on the normalised transcript levels 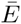 (see Eq.(9)) and *V*, to derive the volume-adjusted transcript uncertainties for the data in Fig. 1

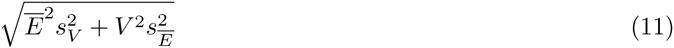

where 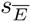 is defined in Eq.(10) and *s*_*V*_ is the SEM for cell volume (raw data provided by Martin Picard). For the case of ETC protein data, since the corresponding experiments in Picard et al. [13] had only a single technical replicate, we derived an uncertainty by simply multiplying the normalised protein value (see Data Normalization) by *s*_*V*_.

### Growth Rate Determination

The speed with which cells proliferate is dependent upon heteroplasmy, as can be seen in Figure S11. However, by day 6 of growth, cell growth appears to change its behaviour, with evidence of saturation; we therefore truncate the raw data to day 5 and calculate the exponential growth rate by linear regression in log-lin space.

### Generative Model Description

We used a Bayesian framework to find the supported parameter values given the data, using the Metropolis-Hastings algorithm [41]. To do this, we included an additional 6 noise parameters, for the features where parameter inference was performed (i.e. all of the features except *N*^+^, which has no free parameters, see Eq.(1)). For these 6 features (*M*_ETC_*, P* ^+^*, M*_gly_*, V, G, R*_max_), we assumed that the data were generated subject to Gaussian noise (see Generative Model Description).

Thus, the full statistical model contains 12 parameters (excluding 6 noise parameters for each feature), with 32 data points which enter the likelihood (after excluding *h*=1 data). To summarise, counting the 6 features which have free parameters, the model consists of 12/6=2 mean parameters per feature, on average. Note that simply fitting linear models to the 6 features in Fig. 1 would also require 2 parameters per feature. The model fit is shown in Fig. 3.

To connect our model of mean cellular behaviour 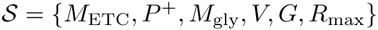, to the data of Picard *et al.* [13], we assume that the sample mean of feature *i* (*y*_*i,j*_) at a discrete value of heteroplasmy *h*=*j* is generated via Gaussian noise 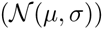 whose mean corresponds to one of the models 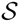,

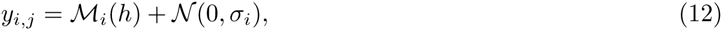

where 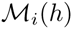 is an element from the set of models 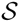. We stress that the data we train our model on, *y*_*i,j*_, is the *sample mean*, rather than the raw data. This is a less common approach; however, we believe that it is appropriate as individual replicates only give us information on the technical variability measured in [13], whereas the total error is a combination of both technical and biological variability. Training our models on individual replicates would be likely to underestimate the true variability of the data, so we favoured training on the sample mean only. This raises the challenge of establishing an appropriately permissive model for our uncertainty in *σ*_*i*_.

We can infer the distribution of the parameters (*θ*) of the models 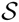, given the data *y*_*i,j*_, using Bayes rule and a prior distribution over *θ* (*P* (*θ*))

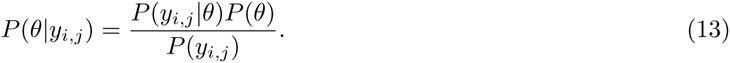

The log-likelihood in this case is

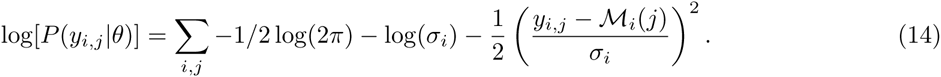

We drop the constant *Σ_i,j_ −* 1/2 log(2*π*) from our log-likelihood, since we will only be interested in differences in the log-likelihood to perform Bayesian inference using the Metropolis-Hastings algorithm [41].

We used exponential priors *σ*_*i*_

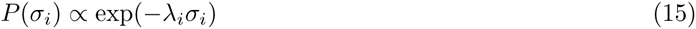

as our error model. The constant *λ*_*i*_ was chosen such that the scale of decay of probability was on the same scale as the range of the data. Noting that 〈*P* (*σ*_*i*_)〉=1/*λ*_*j*_, we chose

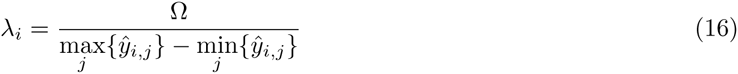

where Ω is a hyper-parameter of the prior and Ω ≥ 0. Note that we may interpret Ω=0 as a uniform prior, since *P* (*σ*_*i*_)=const in this case. In order to make the posterior distribution well-defined, we may think of the case of Ω=0 as a uniform prior *P* (*σ*_*i*_)=unif(0*, α*) for some *α* which is large enough to be never encountered during the finite number of iterations used in our Markov chain Monte Carlo sampling strategy. This is as opposed to an improper uniform prior which would make the posterior distribution unnormalized. In this case *α*=100 is a sufficiently large upper bound to never be encountered in the 10^10^ iterations of the sampler.

We began with Ω=0 as the most permissive choice of prior possible, given the model in Eq.(15). We found that when Ω=0 *the maximum a posteriori estimates were qualitatively similar* to choosing Ω=2 (our final choice which we justify below) see Fig. 3 (Ω=2) and Figure S12A-F (Ω=0). However, we found that the posterior 25-75% confidence intervals supported model fits for *M*_ETC_, *P* ^+^ and *R*_max_ which were relatively poor when Ω=0, compared to Ω=2 (see Figure S12A-F). We determined that large values of *h*^*\**^ were indicative of purely linear fits to the data, which is unlikely given the wider body of evidence demonstrating the nonlinearity of the threshold effect. This is seen in Figure S12G-L (high *h*^*\**^, poorer fit) when compared with Figure S12M-R (low *h*^*\**^, better fit). Comparison between Figure S12G and Figure S12M is particularly noteworthy, where the 25-75% posterior confidence interval for high *h*^*\**^ sub-samples predicts *M*_ETC_ ≈ 0 for all values of *h*, which is physiologically implausible, whereas low *h*^*\**^ sub-samples display non-linear fits which more faithfully track the data. Figure S13 shows that the high *h*^*\**^ mode is of comparable prevalence to the low *h*^*\**^ mode when Ω=0.

We therefore investigated the sensitivity to choice in Ω in Figure S13. We see that increasing Ω reduces the width of the marginal posterior distribution of *h*^*\**^, constraining the posterior distribution to lie around the nonlinear solutions shown in Fig. 3. We found that the permissive prior Ω=2 was sufficient to strongly subdue this, physiologically implausible, large *h*^*\**^ mode. This can be interpreted as a prior belief that our model uncertainty is, on average, 50% of the range of the data (since 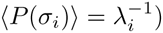. We believe this to be a sensible prior choice, encoding our prior belief that the threshold effect is nonlinear while providing only a gentle constraint on parameters.

We favoured uniform priors on the remaining parameters so that the posterior would be dominated by the likelihood. However, a number of the parameters in the model were uncertain over orders of magnitude; in these cases, we allowed the log of these parameters to take uniform distributions. Explicitly, our priors were chosen as:

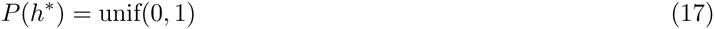

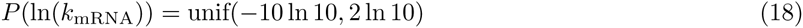

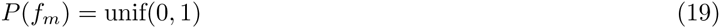

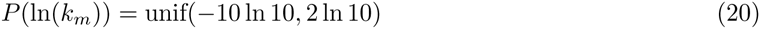

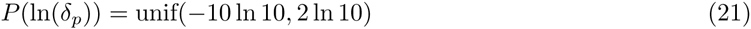

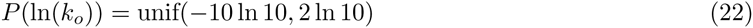

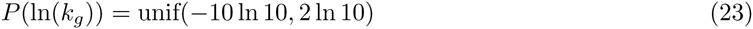

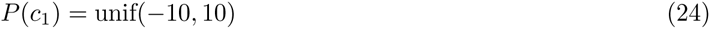

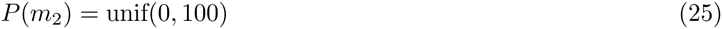

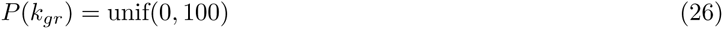

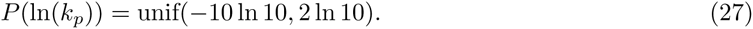

The ranges for *h*^*\**^ and *f*_*m*_ are justified since these quantities can physically only be between 0 and 1. *c*_1_ and *m*_2_ are parameters of linear models for *M*_*gly*_ (see Table S2 and Eq.(5)) for data which has been normalized to the scale of 1; therefore priors were chosen with suitably large ranges. Similarly for *k*_*gr*_, a proportionality constant relating growth to cell volume ((see Table S2 and Eq.(7)), we expect *k*_*gr*_ to be of the order of 1, since the data has been normalized, and chose suitably relaxed priors. The ranges for all other parameters, which were sampled in log-space due to our greater uncertainty of their values, were chosen to be suitably large as to be unlikely to reach the boundary of the prior during sampling with MCMC.

The parameters *β* and *k*_mRNA_ from Eq.(2) were highly correlated. For more efficient chain mixing, we rearranged Eq.(2) into the form

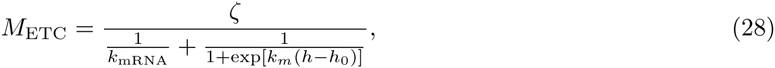

where *ζ*=*β/k*_mRNA_, and used the prior

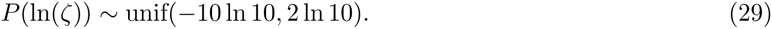

and again used relatively relaxed boundaries for the uniform prior.

We performed the Metropolis-Hastings algorithm [41] to sample from the posterior, using a Gaussian random walk as our transition kernel, whose covariance matrix was determined from a trial run of the adaptive Metropolis algorithm [42]. All code was written in either Python or C, and is available upon request. The MCMC chain trajectory is presented in Figure S2.

## Supporting Information

### Text S1

#### Justification of ETC mRNA and protein

Consider a single molecule of wild-type mtDNA which, when transcribed, generates mRNA for the electron transport chain (ETC), which we denote as *m*_*ETC*_. Transcripts are generated according to a deterministic process (stochasticity in gene expression [43] is neglected in this picture) with rate (*β*) and also passively degrade at some basal rate 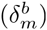. We consider a controlled, active degradation process 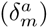 that acts in addition to the background level. Thus, at the single mtDNA level, we may write down the differential equation

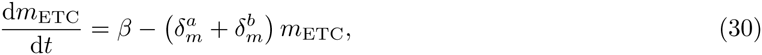

where we assume that *β*, 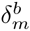 are constant.

Control of the expression levels of different mitochondrial genes is manifest at the level of mRNA degradation [25], because mtDNA is transcribed as a single polycistronic transcript [24]. We therefore use the simplifying assumption that, in the pathogenic case, mitochondrial mRNA is also controlled at the level of degradation. Thus we allow the active degradation to vary with heteroplasmy 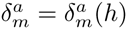, and assume the transcription rate to be constant.

Cells are measured at steady-state, so setting the derivative to zero yields

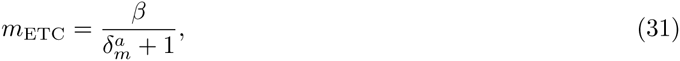

where 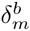 has been absorbed into the definitions of *β* and 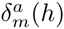. Assuming *N* ^+^ scaling at the cellular level yields the expression

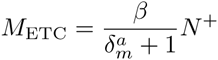

for ETC mRNA at the cellular level. Dropping the *a* superscript yields Eq.(2).

Note that, in our Bayesian inference, we chose to express the constant *h*_0_ in Eq.(3) in terms of the critical heteroplasmy *h*^*\**^ using the expression *h*_0_=*h*^*\**^ − ln[(1 − *f*_*m*_)/*f*_*m*_]/*k*_*m*_, where 0 < *f*_*m*_ ≤ 1. This simply expresses the location of *h*^*\**^ in terms of the fraction *f*_*m*_ of the sigmoid’s maximal value. Intuitively, if *f*_*m*_ and *k*_*m*_ are sufficiently large, *h*^*\**^ signals the beginning of reduction in ETC degradation.

For ETC protein, we assume that the following equation holds at the cellular level

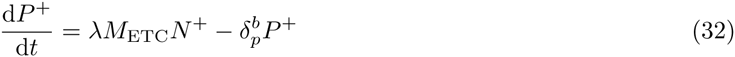

where *λ*=const, and we assume there is no active degradation of ETC protein. At steady-state

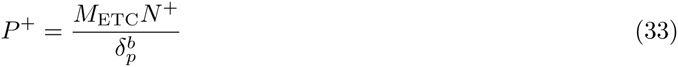

where *λ* is absorbed into the definition of 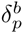. Dropping the *b* superscript yields Eq. (4).

#### Justification of cell volume scaling for energy demand

A reasonable general model for the way in which energy demands of a mammalian cell scale with its volume (*V*) is *k*_1_*V* + *k*_2_*V* ^2/3^ + *K* + *f* (*N, V*), where *k*_*i*_ are proportionality constants. Each term may be interpreted as: *k*_1_*V* are demands which scale with cell volume; *k*_2_*V* ^2/3^ scale with cell surface area; *K* are demands which are constant for a cell (for example, the cost of replicating the genome); and *f* (*N, V*) is an unknown function corresponding to proton leak, which potentially depends upon mitochondrial mass (or alternatively mtDNA copy number, *N*) and cell volume. These are the dominant energy demands of mammalian cells, as determined by [21–23].

Many energy-consuming processes in mammalian cells directly depend upon cell size; for example, a model system used by Buttgereit *et al*. found that ~30% of oxygen consumption corresponded to plasma membrane transporters, and ~20% corresponded to protein synthesis [21]. We assume that protein synthesis scales proportionally with *V*, because 60% of total cellular dry mass is protein [44] and mass scales with volume. Also, we may assume that plasma membrane energy consumption scales with cell surface area, which scales with *V* ^2/3^. By using volume and surface area contributions alone, we may account for ~50% of the energy demands of the cell, which is 63% of the accountable energy demands for this model system since only 80% of total respiration rate could be attributed to particular processes in their study [21].

Thus, assuming that all energy consumption is due to surface area or volume contributions then, using the data of [21], a reasonable model for energy demand might be *f* (*V*)=0.4*V* + 0.6*V* ^2/3^, the volume parameter being 20/(20 + 30)=0.4. However, we see in Fig. 1D that the normalized volume data lie in the range 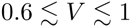. In this region, the functions *f* (*V*) and *g*(*V*)=*V* are similar, with a difference of no more than ~7%. Thus, *g*(*V*)=*V* is a reasonable approximation for total energy demands in this case.

We note, however, that the above proportions depend on the environment of the cell [21, 22] as well as the tissue type (reviewed in [23]), often showing variation on the order of tens of percent. In light of this uncertainty, and for the sake of parsimony, we make the simplifying assumption that total energy demand scales purely with cell volume, see Eq.(6).

#### Expected cell volume and growth rate

Two of the simplest models for how cells may grow throughout the cell cycle are linear and exponential growth. We show below that a relationship exists between growth rate and the mean cell volume in an asynchronous population of cells under a linear model. Furthermore, assuming an exponential model, growth rate and mean cell volume are independent.

Firstly, we assume that the number of cells obey a pure-birth process, in other words the death rate of cells is negligible. If the initial number of cells (*N*_0_) is large, then we can use a deterministic model of cell growth, *N* (*t*)=*N*_0_ exp(*Gt*), where *N* (*t*) is the number of cells at time *t* and *G* is the growth rate of cells, as described in Eq.(7). Assuming that the number of cells doubles every cell cycle period (*t*_*d*_), then

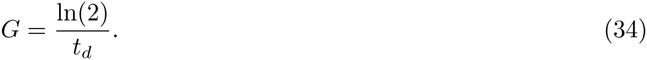

Under a linear model of cytoplasmic growth through the cell cycle, the volume of an individual cell may be written as

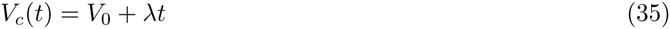

where *V*_*c*_ is the volume of an individual cell, *V*_0_ is the volume of a cell just after division (assumed to be constant for all cells) and *λ*=*V*_0_/*t*_*d*_. In an asynchronous population, we assume that each cell is distributed uniformly through the cell cycle, in other words

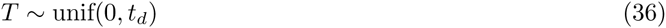

where *T* is a random variable describing the position in time, of a cell in its cell cycle.

We wish to find the expected value of cell volume, (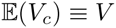, as described in Eq.(6)), given the assumption of Eq.(36). Eq.(35) can be viewed as a transformation of the random variable *T*. If *X* is a continuous random variable, then for any transformation 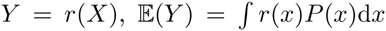, where *P*(*x*) is the probability distribution corresponding to the random variable *X* [45]. This implies that 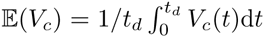. For Eq.(35), this yields 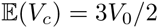, but since *V*_0_=*λt*_*d*_, then Eq.(34) yields

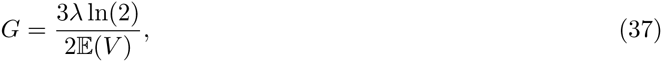

i.e. 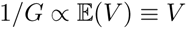.

If, however, we assume an exponential model of cell growth through the cell cycle

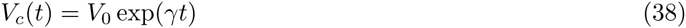

then *t*_*d*_=ln(2)/γ and 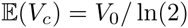 which cannot be written in terms of *t*_*d*_ and therefore *G* is independent of *V*.

The above makes intuitive sense: if a cell grows linearly, then a larger cell will need more time to double in size than a smaller cell, if their growth rates are the same. On the other hand, if a cell grows exponentially, then regardless of its initial size, the doubling time is constant, given a fixed cytoplasmic growth rate.

Since there is presumably a wide class of cell growth dynamics where cell size is dependent on growth rate, we favoured a linear model for its simplicity. Measurements by Tzur *et al*. show that, on average, under both a linear and exponential model of cytoplasmic volume growth, the rate constant varies with time. [46] However, the implication of this for the relationship between *V* and *G* remains unclear.

### Text S2

Below, we will propose potential experiments to test the corresponding claims made in Key Claims and Predictions of Biophysical Model of Heteroplasmy.

#### Wild-type mtDNA density homeostasis is maintained until a minimum volume is reached at the critical heteroplasmy

If *N* ^+^/*V* is a quantity kept under homeostasis, then under wild-type conditions, perturbations to mtDNA copy number may be expected to cause changes in cell volume. This might be testable by reducing mtDNA copy number with chemicals such as ddC, or increasing it through PGC-1*α* overexpression, which, in the absence of other homeostatic effects, we expect to reduce and increase mean cell volume respectively.

A second testable prediction is that a minimum cell volume (*V*_min_) causes bioenergetic toggling at *h*^*\**^. We should be aware of the two potential interpretations of *V*_min_ raised: (1) bioenergetic and (2) mechanical. If *V*_min_ is bioenergetic, then raising the power demands of the cell which do not scale with volume, may induce *h*^*\**^ to be encountered earlier. This could be achieved by increasing the amount of DNA in the nucleus which must be replicated, creating a one-time cost to the cell per cell cycle. This is not expected to affect any mechanical constraints since DNA content is not directly indicative of nuclear size [47]. Ideally, the amount of DNA introduced should be large (i.e. billions of base pairs), be replicated, and not interfere with normal functioning of the nucleus. This could be achieved by chemically inducing polyploidy, for instance by using Noscapine [48]. The location of *h*^*\**^ could again be determined by performing RNA-seq, and observing the upregulation of glycolysis with heteroplasmy.

If increasing the energy demands of the nucleus yields no change in the distribution of *h*^*\**^, then a mechanical constraint could be more relevant. This could be tested by perturbing cell volume. Reducing cell volume under this hypothesis is expected to shift *h*^*\**^ to lower values of heteroplasmy, which could be determined via RNA-seq.

#### Mutant mtDNAs do not contribute to the mitochondrial mRNA pool

If mutant mRNAs have a transcription defect, then the abundance of mutant tRNAs relative to wild-type mtDNAs would be expected to be smaller than *h*. This measurement could be performed by qPCR, to probe mutant and wild-type tRNA copy numbers.

#### Mitochondrial tRNAs are relatively localised to their parent mtDNA

The spatial distribution of mitochondrial tRNAs relative to mtDNA would be most directly determined by fluorescent labelling of mitochondrial tRNA and mtDNA. MtDNA labelling could be achieved through picoGreen staining [49]. Labelling of processed tRNAs within mitochondria is more difficult, but methods exist for labelling mRNA in both fixed cells [50] and dynamically [26, 51] within mitochondria, which may be informative.

#### Cell volume is not explained by cell cycle variations

To separate the potential confounding influence of the cell cycle on mean cell size, heteroplasmic cells could be transfected with Fucci markers [52], and relative enrichment of cell cycle stages determined.

#### Cells proliferate inversely with their size

To determine the dependence of growth rate on mean cell volume, wild-type cells could be synchronised, and sorted by their volume. These cells could then be plated and released from synchronisation, and the growth rate of cells measured similar to that described in Materials and Methods. Synchronisation is necessary, because cell volume is expected to vary by a factor of 2 through the cell cycle, so any sorting would otherwise be strongly confounded by the cell cycle. A potential alternative to synchronisation, which can be stressful to cells, is to label genes associated with a particular stage of the cell cycle, and sort based on both this fluorescence signal and cell volume.

#### Maximum respiratory capacity linearly tracks ETC protein content

Measurements of maximum respiratory capacity at *h*=0, as well as measurement of ETC protein levels at *h*=0.6, may help determine whether a simple linear relationship is sufficient, or whether a more complex model is justified.

#### Summary

A summary of the experimental proposals outlined are given in Table S1

### Text S3

#### Relative OXPHOS contribution to energy supply

It is interesting to observe the relative contributions of oxidative phosphorylation and glycolysis to power supply. Since Eq.(6) states that energy supply=demand, where demand corresponds to cell volume, the ratio *f*_*o*_=*k_o_P* ^+^/*V* determines the relative contribution of OXPHOS to energy supply, see Figure S14.

For *h < h*^*\**^, we see that OXPHOS has decreasing contributions to energy power supply. At *h*^*\**^, OXPHOS contributions stabilize with 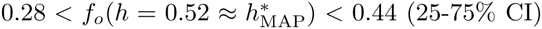. The heteroplasmy at which OXPHOS contributions are stabilized corresponds to the hypothesized demand/supply toggle, where the cell attempts to increase energy supply as opposed to reducing energy demand.

The value of *f*_*o*_ where OXPHOS contributions become stabilized (*f*_*o*_(*h*^*\**^)) may have wider significance. Mitochondrial metabolism, and especially mitochondrial membrane potential, is connected to a variety of biosynthetic pathways [1] and crucial for maintaining cellular proliferation [39]. *f*_*o*_(*h*^*\**^) may represent a minimum ETC flux, relative to energy demand, for mitochondria to support their mitochondrial membrane potential without the aid of glycolytic ATP. Below *f*_*o*_(*h*^*\**^), we might predict that cells run ATP synthase in reverse, hydrolysing glycolytic ATP to maintain membrane potential.

### Text S4

#### Alternative models: Mutant mtDNA and transcription

Eq.(2) states that mutant mtDNAs do not contribute significantly to the transcript pool. We can relax this constraint by replacing Eq.(2) with

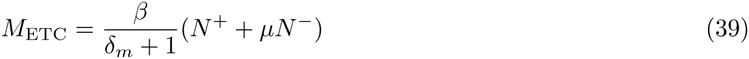

where 0 ≤ *μ* ≤ 1 and *N*^*−*^=*hN*, where *N* is the total number of mtDNAs, which we treat as a constant.

Using the uniform prior

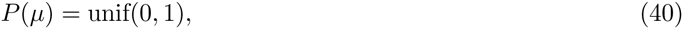

we sampled from the posterior, as described in Materials and Methods. The MCMC trajectory is shown in Figure S3. The marginal posterior density for *μ* in Figure S3, shows that *μ* ≈ 1 is the most likely value of the parameter, in other words mutant mtDNAs contribute equally to the transcript pool, compared to wild-type molecules. However, observing the model fit for *M*_ETC_ in Figure S5 and Figure S15, it is clear that this model favours a linear fit to the data, with large uncertainty.

#### Alternative models: tRNA misincorporation model

Eq.(4) states that ETC protein is generated when ETC mRNA is in contact with wild-type mtDNA, suggesting that tRNAs affected by the MELAS mutation, leucine-UUR, remain local to their parent mtDNAs. The alternative is that mitochondrial tRNAs are well diffused amongst mitochondrial mRNAs. If we assume that a mutant tRNA causes a misincorporation during translation with 100% efficiency, then the number of misincorporations per protein follows a binomial distribution. We assume that the probability of a single misincorporation is *h*. We further assume that proteins have a mutational tolerance of *x* misincorporations, or less, before they are considered mutated (and consequently degraded). With these assumptions, the expected proportion of mutant proteins (*m*_*p*_) will be

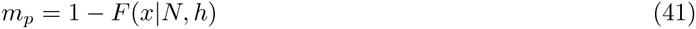

where *N* is the number of leucine-UUR residues per protein, and *F* (*x|N, h*)=*P* (*X* ≤ *x*) is the cumulative distribution function of the binomial distribution, for *N* trials, *x* successes, and probability of success *h*. A plot of *m*_*p*_ is given in Figure S6, for different mutational tolerances *x* against heteroplasmy.

We can therefore use an analogous expression to Eq.(4) for *P* ^+^, in the case of well-diffused tRNAs

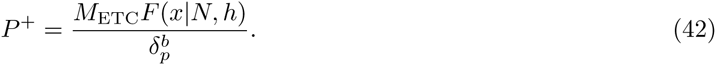

By replacing Eq.(4) with Eq.(42), we again sampled from the posterior as described in Materials and Methods. We chose *N*=8, which is the average number of susceptible residues in the 11 mitochondrially-encoded subunits considered (see caption of Fig. 1) [9]. Our prior for the unknown tolerance to misincorporations, *x*, was chosen as a discrete uniform prior

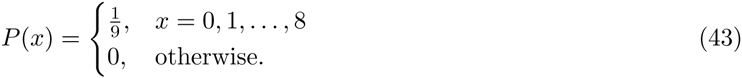

The MCMC trajectory is shown in Figure S4, and the model fit in Figure S6. The model fit shows that, whilst the maximum a posteriori estimate for *M*_ETC_ fits the data more closely, the 25-75% confidence interval is poorer than the model presented in the main text (see Fig. 3 and Figure S15). All other features are comparable. Furthermore, observing the marginal posterior distribution of the misincorporation tolerance *x* in Figure S4, we see that the most likely value of the parameter is *x*=*N*=8. In other words, ETC proteins are immune to the MELAS mutation, which we believe to be incorrect [9].

### Text S5

#### Transformation to Per-Cell Dimensions using Cell Volume

In this section we show that it is necessary to multiply measurements of protein and mRNA levels, when determined by Western blot and RNA-seq respectively, by cell volume to transform the data to mean cellular measurements.

Consider a Western blot experiment determining the levels of a gene (gene *i*) in two conditions (A and B). Denote the number of proteins per cell of gene *i*, as 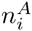, where the superscript denotes condition A. Let us also denote the total number of proteins per cell as *N*^*A*^.

When one performs a Western blot, the protein of interest is stained with an antibody, cells are lysed, and a sample of fixed protein mass (*m*) is taken from the lysate. If we denote the number of proteins for the gene of interest in the sample as 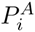, then we may write

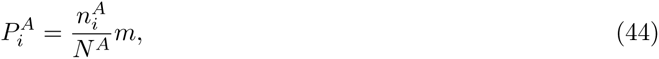

since the proportion of protein *i* in the sample is determined by the proportion of protein *i* in the proteome 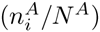. Western blot experiments also tend to be normalised by a loading control (*c*), so the normalised measurement we have access to is

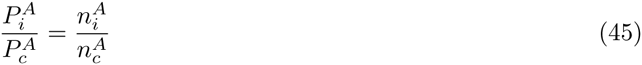

which corresponds to the data given in Picard et al. [13].

Now, consider a perturbation in condition B, causing the amount of protein for gene *i* to be 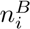, and the mean cell volume to experience a fold-change *V*_*f*_, as in Figure S10. Consequently, *N* ^*B*^=*V*_*f*_*N*^*A*^, since total protein content scales with the volume of the cell. Using the reasonable assumption that the loading control is a gene whose expression also scales with cell volume (e.g. *β*-actin, as in Picard et al. [13]), then 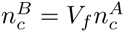. It follows that

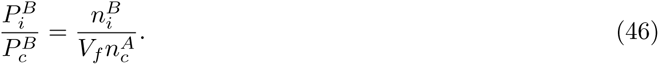

Then, if we are interested in the relative fold-change expression of the protein between the two conditions, then we take the ratio

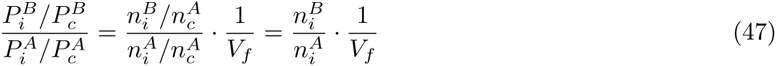

Thus, the quantity on the left hand side of Eq. (47), which is what one usually measures in a Western blot, has a multiplicative-bias of 1/*V*_*f*_. Therefore, if one is interested in per-cell protein changes, the appropriate quantity of interest is

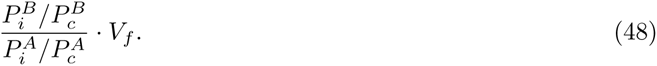

Hence, we multiply each protein measurement by 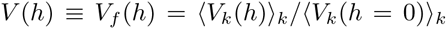, where 〈…〉_*k*_ denotes the sample mean over technical replicates *k*, such that *V* (0)=1.

A similar argument applies to RNA-seq data, since a fixed mass of mRNA is extracted for an RNA-seq experiment, so an analogous pair of equations to Eq.(44) in conditions A and B holds. Using the assumption that *N* ^*B*^=*V*_*f*_ *N* ^*A*^, and denoting the number of mRNA molecules in each sample with *M*, it can be shown that

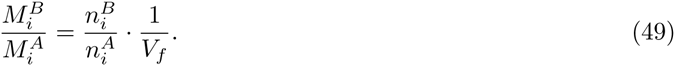

**Figure S1.**
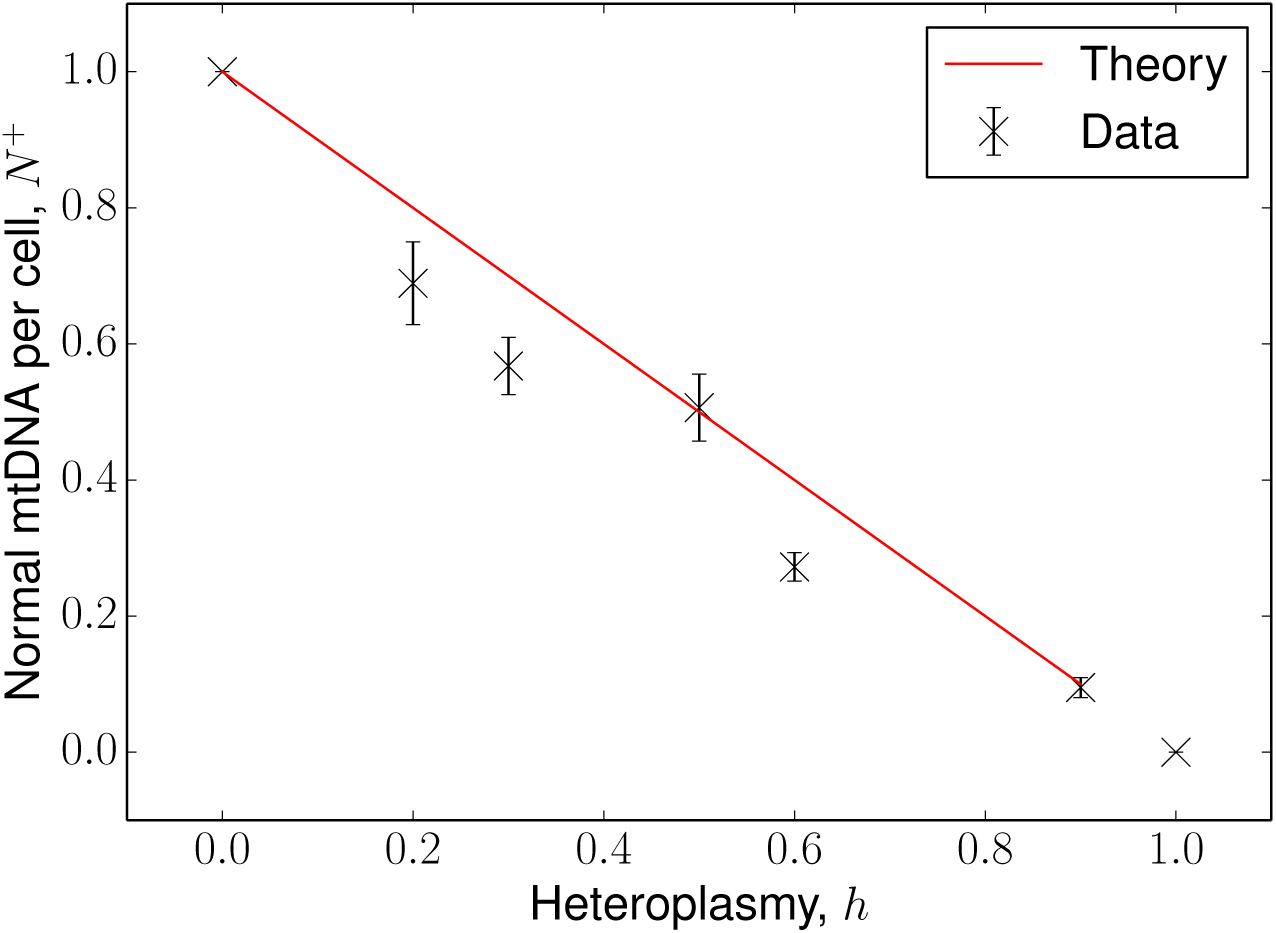
Model fit for wild-type mtDNA copy number. Data for mtDNA copy number from Picard *et al*. [13] was multiplied by (1-h), as was the SEM. Displaying the model *N*^+^=*N*(1 − *h*), for *N*=const=1.

**Figure S2.**
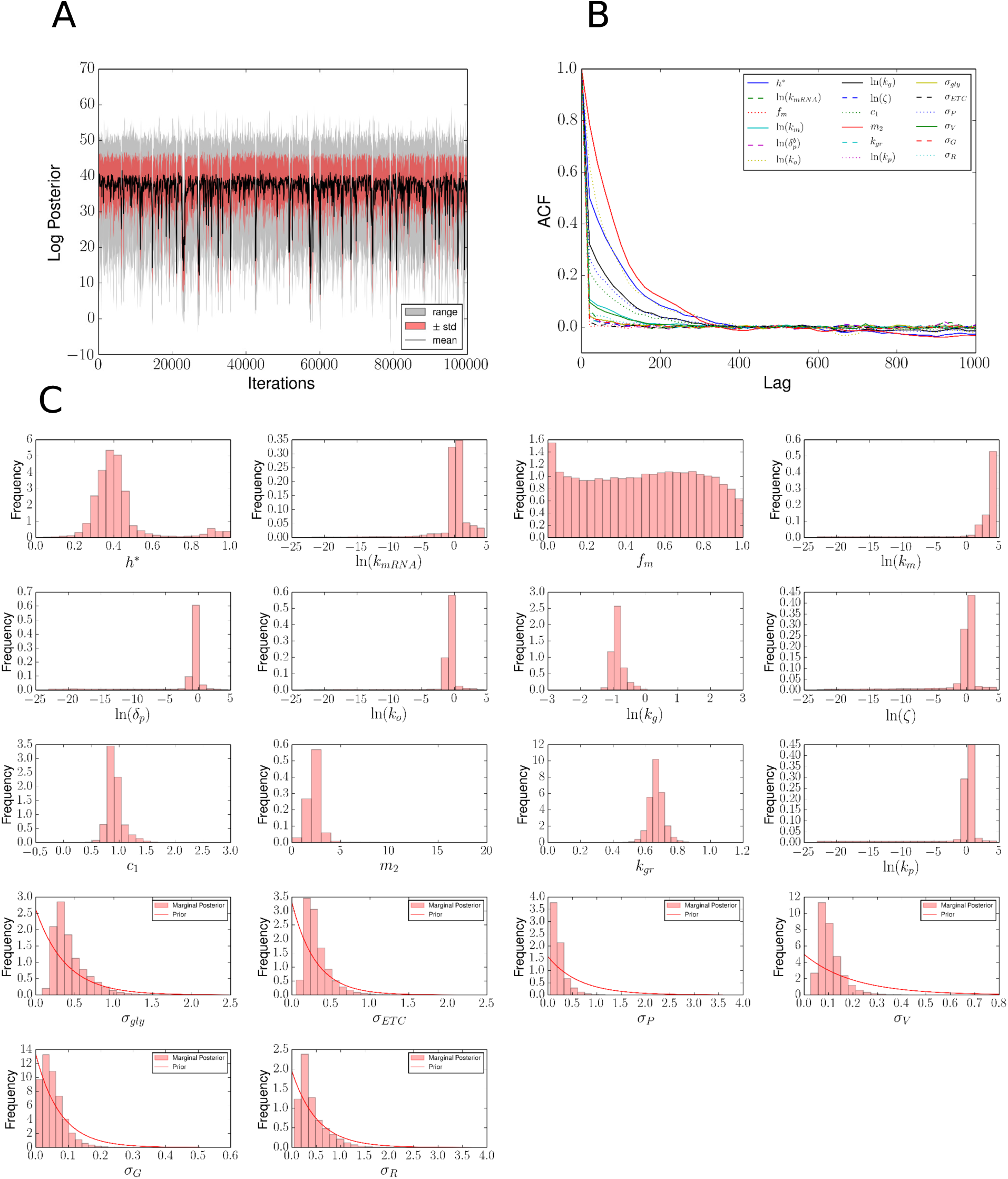
Metropolis-Hastings posterior samples for main model. 10^10^ iterations of Metropolis-Hastings were performed, which were thinned to 10^5^ samples. The hyperparameter for model uncertainty, Ω, was chosen as Ω=2 (see Materials and Methods). A. Trajectory of the unnormalised log posterior after thinning. Samples are split into bins of 100 iterations; displaying mean, standard deviation and range of each bin. B. Autocorrelation function for each parameter, on thinned samples. C. Approximate marginal posterior distributions for each parameter in the model. Exponential priors for model uncertainties are plotted, all other priors are uniform (see Materials and Methods).

**Figure S3.**
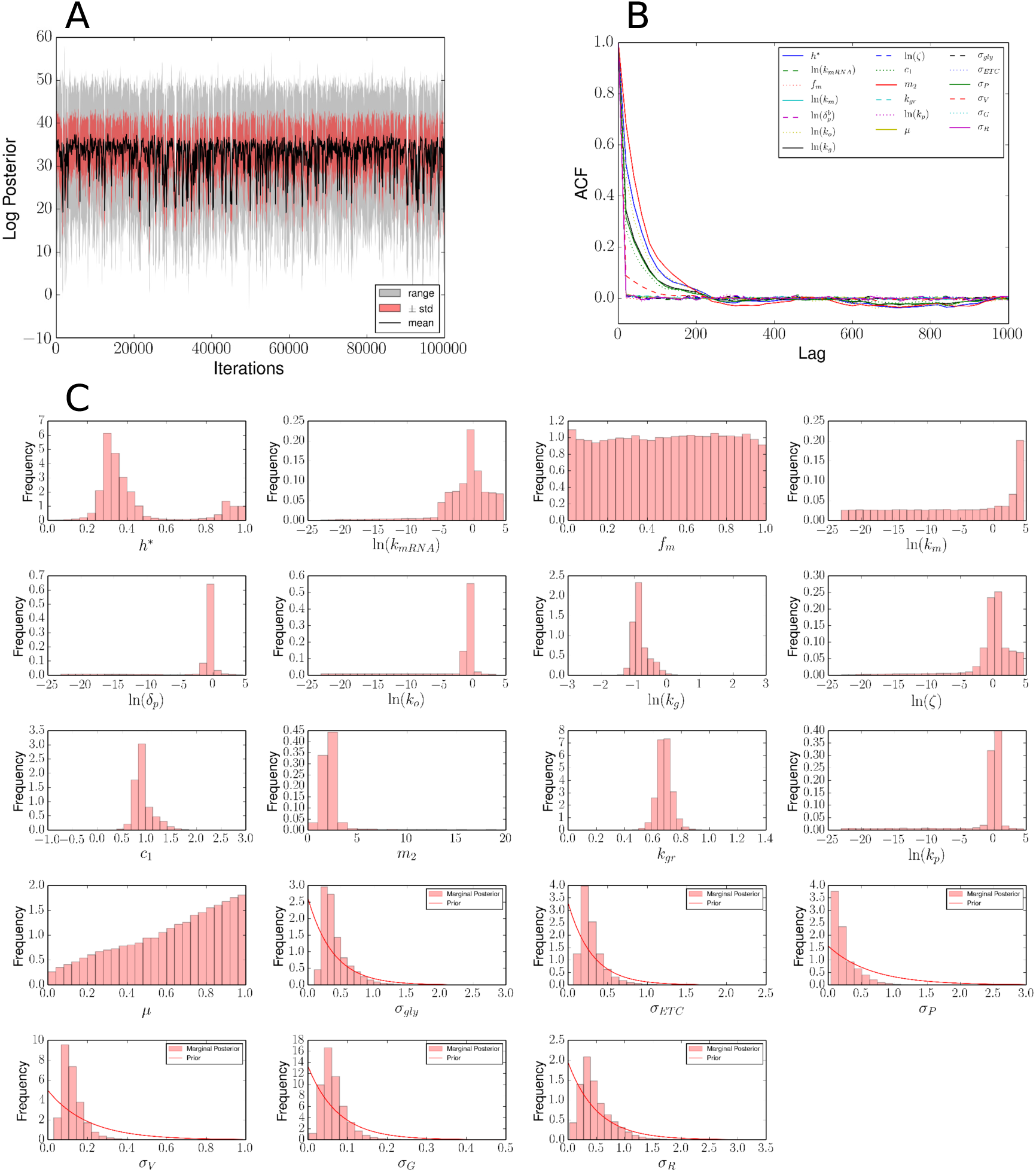
Metropolis-Hastings posterior samples for mutant transcription model. 10^10^ iterations of Metropolis-Hastings were performed, which were thinned to 10^5^ samples. The hyperparameter for model uncertainty, Ω, was chosen as Ω=2 (see Materials and Methods). The prior for the additional parameter *μ* was chosen to be a uniform distribution between 0 and 1. A. Trajectory of the unnormalised log posterior after thinning. Samples are split into bins of 100 iterations; displaying mean, standard deviation and range of each bin. B. Autocorrelation function for each parameter, on thinned samples. C. Approximate marginal posterior distributions for each parameter in the model. Exponential priors for model uncertainties are plotted, all other priors are uniform (see Materials and Methods). Note the marginal distribution of *μ* peaks near 1.

**Figure S4.**
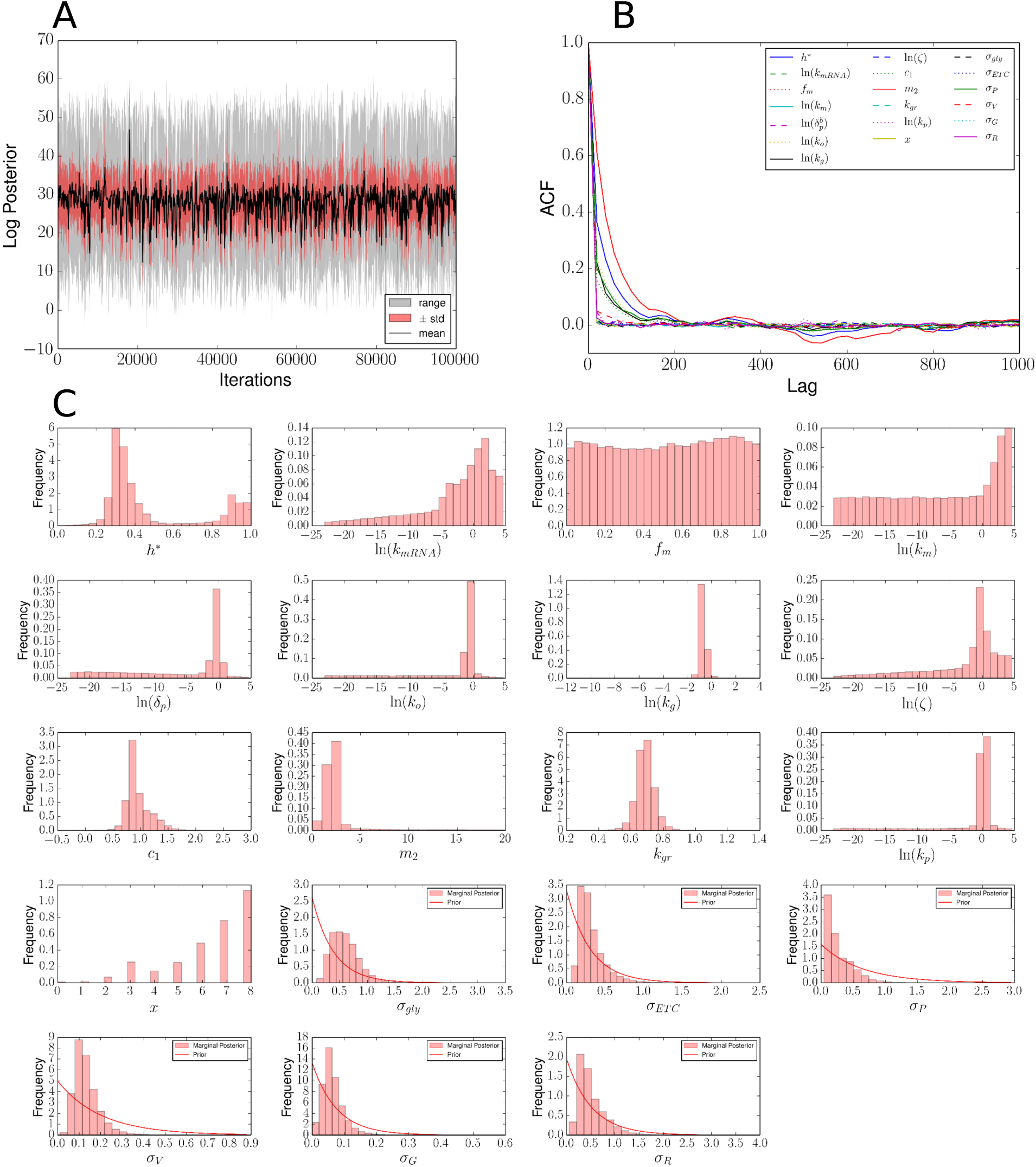
Metropolis-Hastings posterior samples for tRNA misincorporation model. 10^10^ iterations of Metropolis-Hastings were performed, which were thinned to 10^5^ samples. The hyperparameter for model uncertainty, Ω, was chosen as Ω=2 (see Materials and Methods). The prior for the additional parameter *x* was chosen to be a uniform discrete distribution between 0 and 8. A. Trajectory of the unnormalised log posterior after thinning. B. Autocorrelation function for each parameter, on thinned samples. C. Approximate marginal posterior distributions for each parameter in the model. Exponential priors for model uncertainties are plotted, all other priors are uniform (see Materials and Methods). Note the marginal distribution of *x* peaks at 8, which we expect to be incorrect.

**Figure S5.**
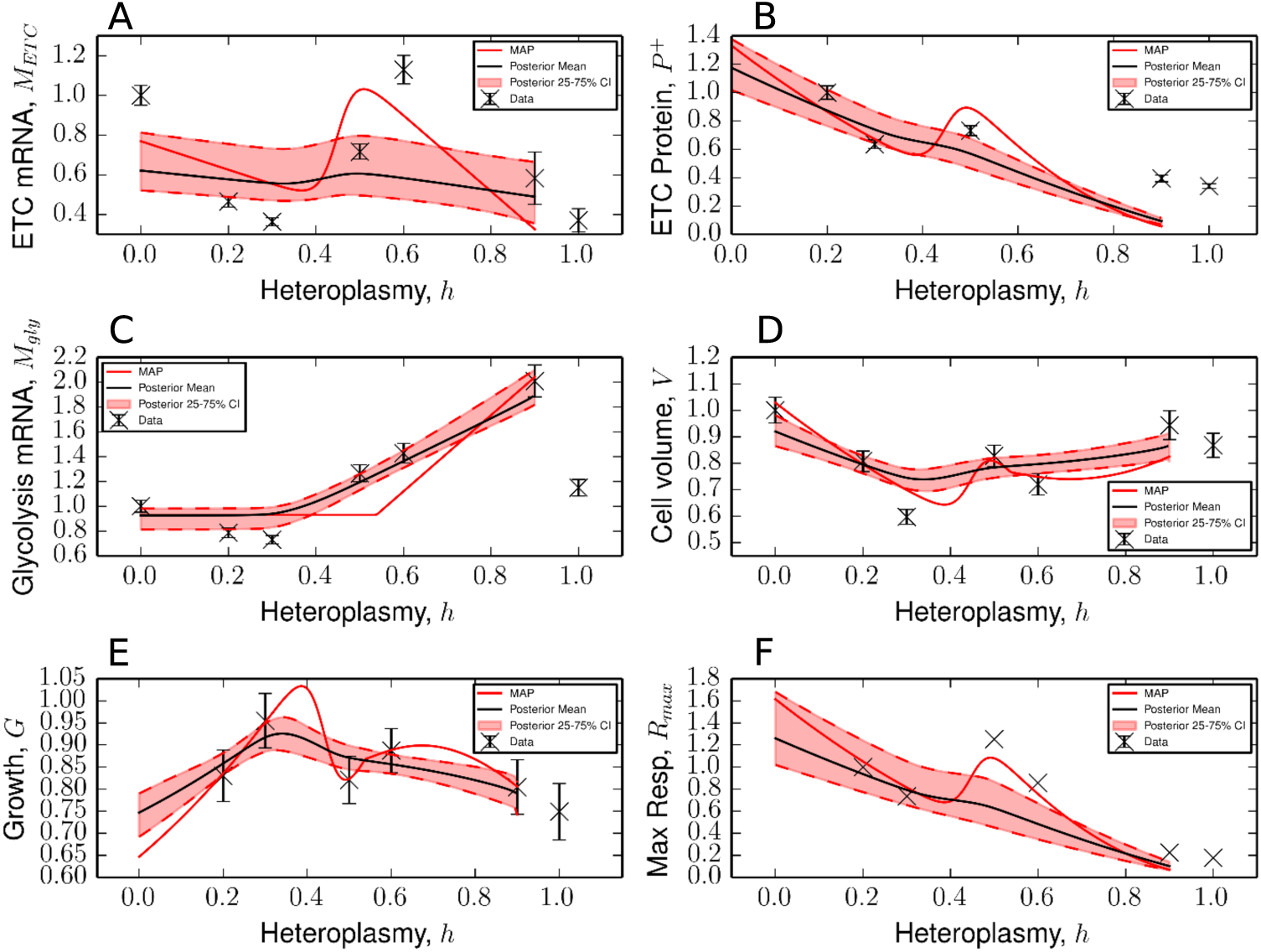
Alternative mutant transcription model. A-F. Model fit when Eq.(2) is replaced with Eq.(39). The 25-75% CI is flatter when compared with Fig. 3. We find this is due to the model more frequently selecting linear fits to the data, see Figure S15.

**Figure S6.**
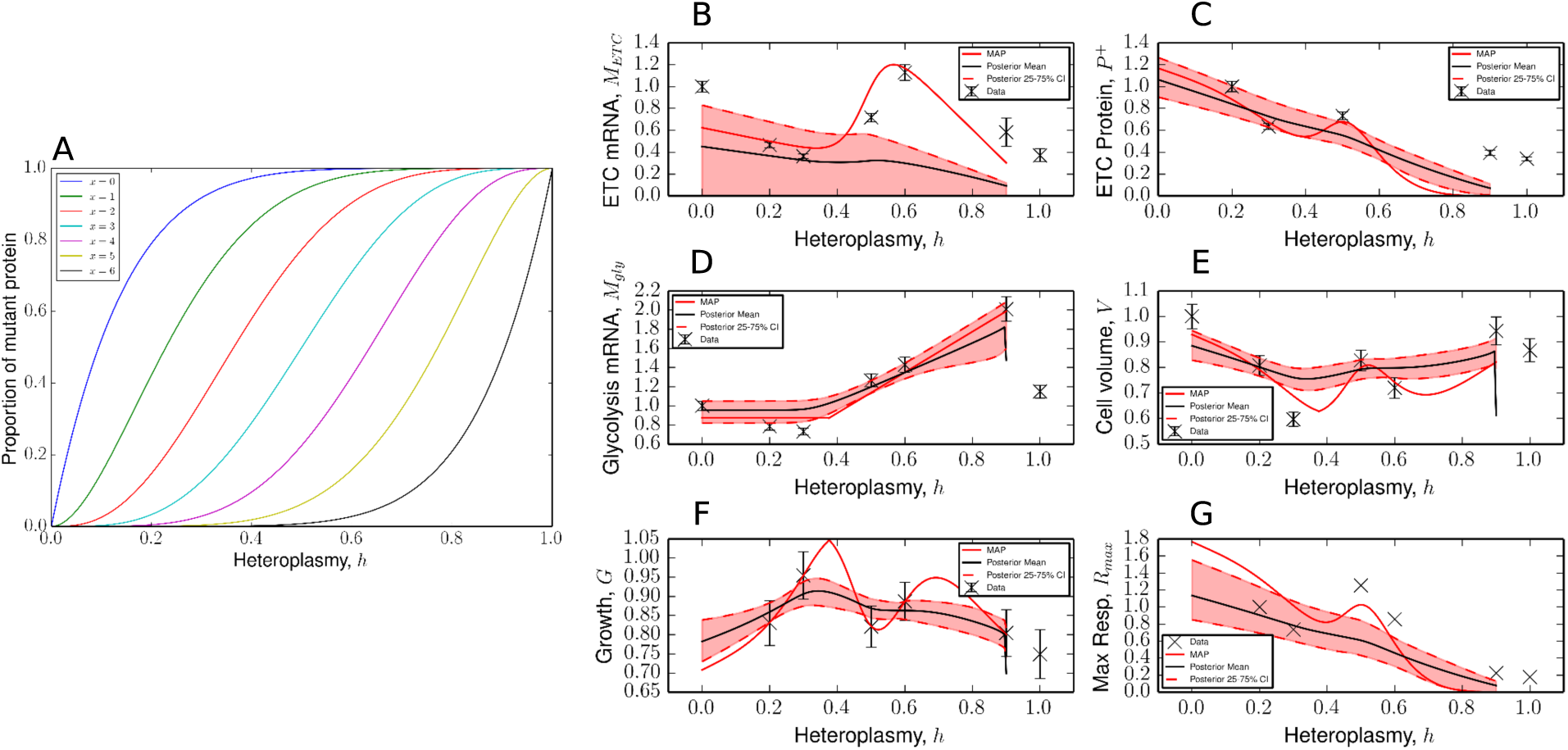
Alternative tRNA misincorporation model. A. Expected proportion of mutant protein due to the MELAS mutation, given that each protein can tolerate *x* mutated residues. The chain length used is *N*=8, which is the mean number of susceptible residues across all mitochondrially-encoded peptides, excluding ATP8 and ATP6 [9]. B-G. Model fit when Eq.(4) is replaced with Eq.(42). *M*_ETC_ is qualitatively fitted more poorly than Fig. 3 (see also Figure S15), although the maximum a posteriori estimate is a closer fit, when compared with Fig. 3.

**Figure S7.**
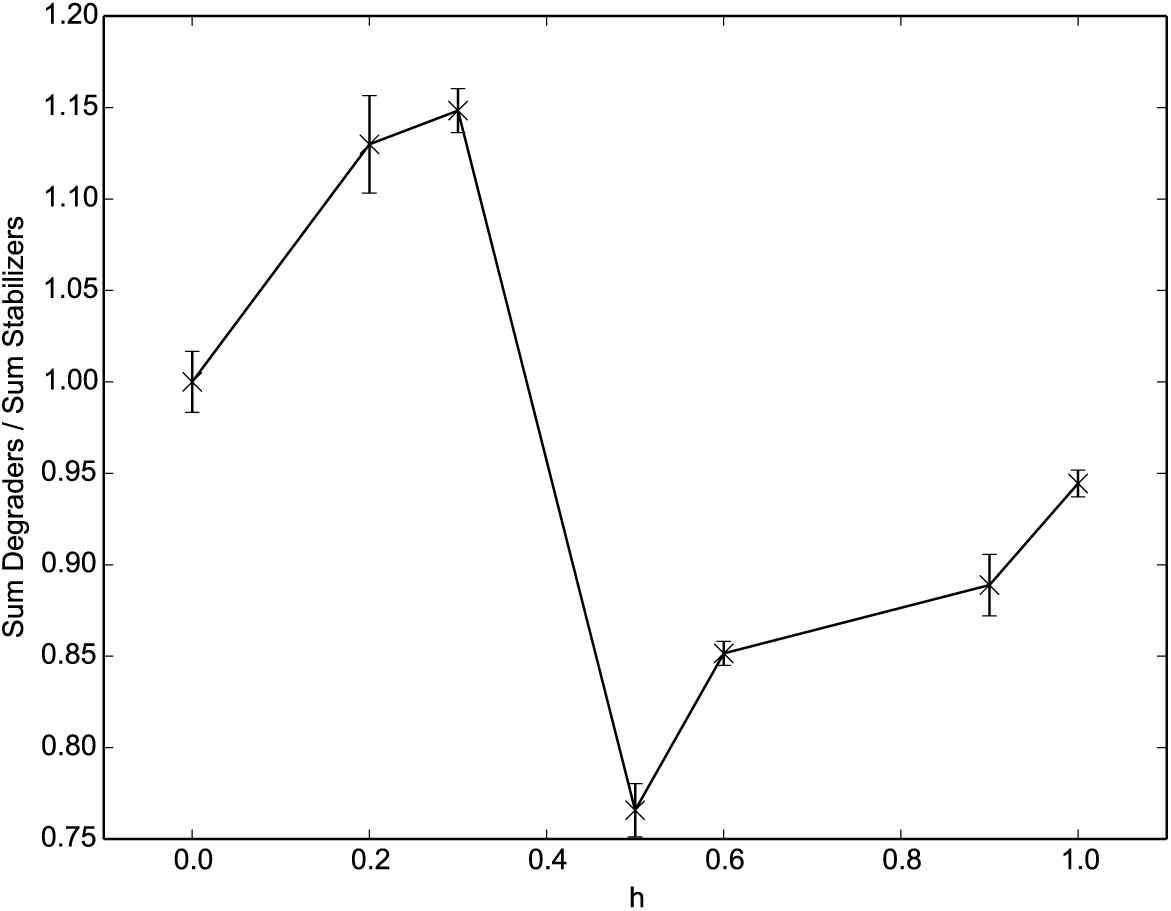
Corroborating evidence for ETC mRNA degradation from Picard *et al*. Ratio of mitochondrial mRNA degraders (PDE12, PNPT1, SUPV3L1) to stabilizers (MTPAP, LRPPRC, SLIRP), see Ref. [25] and references therein. We observe qualitative similarity in 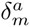 (see Fig. 6) and the ratio of normalised genes, both showing strong downregulation between *h*=0.3 and *h*=0.5. The numerator and denominator were normalised according to Eq.(9). Errors result from error propagation of a ratio, where the error for the numerator and denominator are derived using Eq.(10).

**Figure S8.**
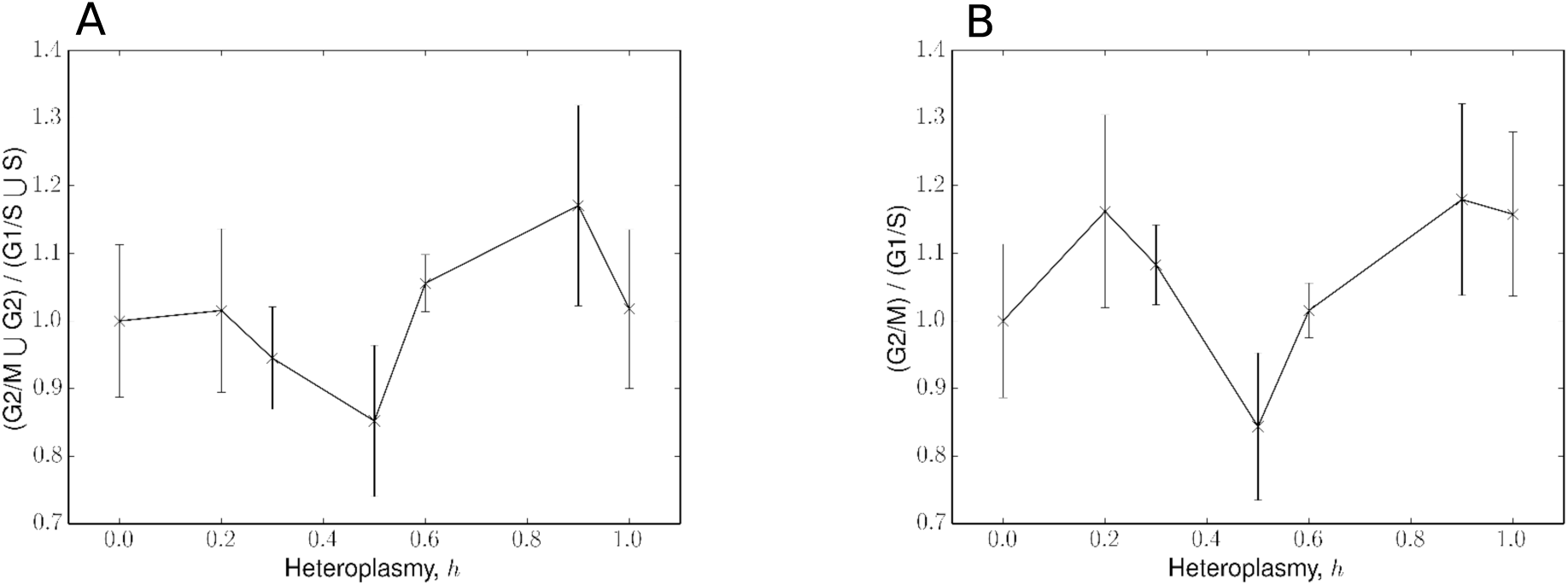
Variation of cell cycle markers with heteroplasmy. A list of cell cycle markers, taken from [28] were normalised according to Eq.(9), yielding gene lists for G1/S, S, G2 and G2/M phases. A. The ratio of G2/M to G1/S genes yielded no obvious trend with heteroplasmy. B. G2/M and G2 gene lists were combined (G2/M ⋃ G2), as were G1/S and S (G1/S ⋃S). This again yielded no obvious trend with heteroplasmy. Errors result from error propagation of a ratio, where the error for the numerator and denominator are derived using Eq.(10).

**Figure S9.**
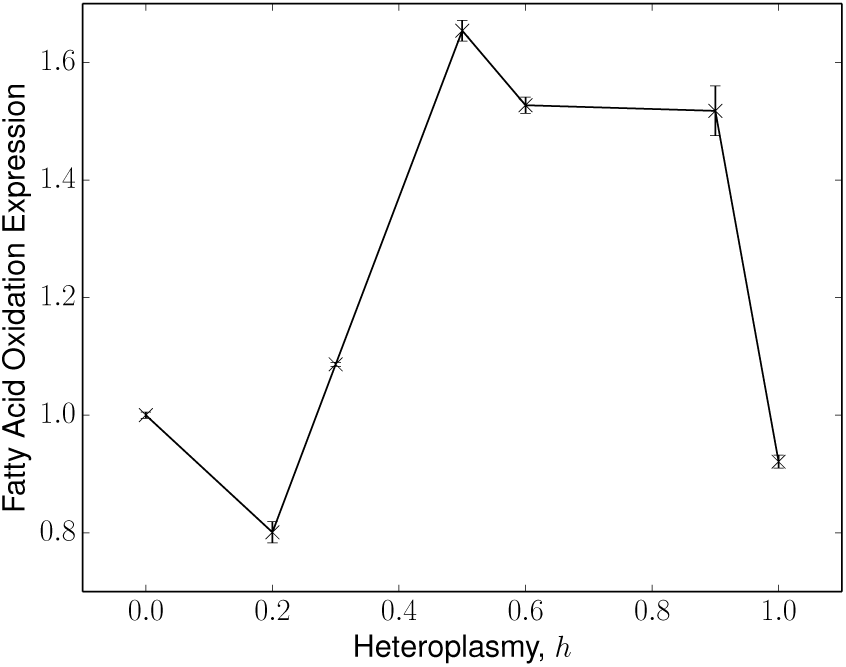
Expression of fatty acid oxidation enzymes with heteroplasmy. Showing variation of the genes (ACADVL, ECHS1, HADH and ACAA2) with heteroplasmy, normalised according to Eq.(9). It can be seen that these metabolites are downregulated between *h*=0.9 → 1, so fatty acid oxidation does not appear to be supporting the maintained cell volume and growth rates, over this range of heteroplasmy. Errors are calculated using Eq.(10).

**Figure S10.**
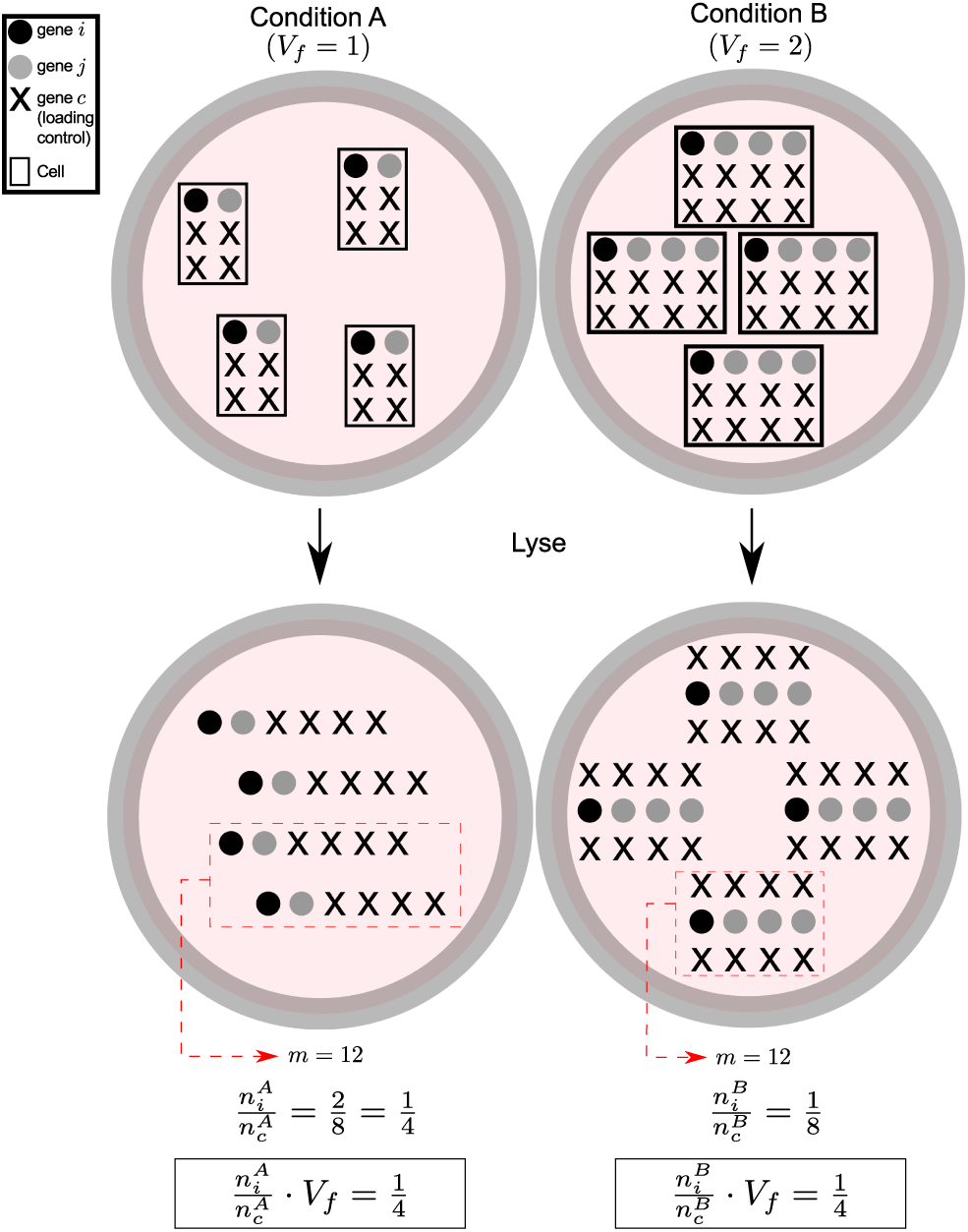
Transformation of Western blot (or RNA-seq) data to per-cell dimensions. Consider a Western blot experiment, where we are interested in the fold-change expression of gene *i* per cell (*n*_*i*_), and cell volume has a fold change *V*_*f*_=2 between conditions A and B. In this example, 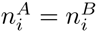. Taking an unbiased sample of size *m*=12 from each condition, and dividing by the loading control, yields a quantity 1/*V*_*f*_ too small. It is necessary to multiply the ratio by *V*_*f*_, to get an accurate measurement of gene *i*, in the context of strongly varying cell volume, as is the case in Picard *et al.* [13]. A similar argument holds for RNA-seq data, which also uses a fixed mass of RNA as the starting sample.

**Figure S11.**
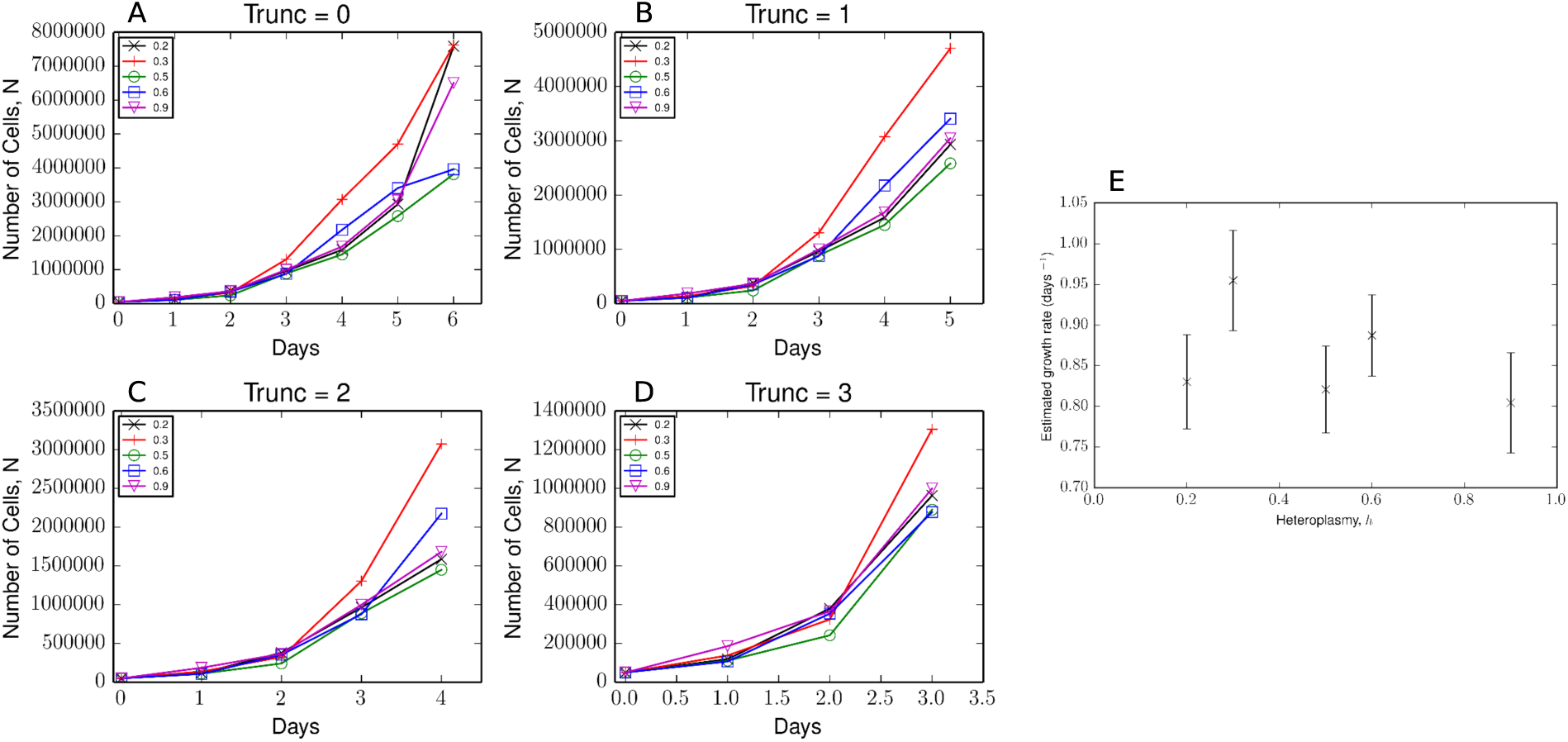
Cell proliferation data from [13]. A-D. Number of cells (*N*) versus number of days of incubation, for different heteroplasmies, where a number of data points have been truncated (Trunc) from the right. Growth appears to be non-exponential by day 6, and is therefore removed subsequently. E. Slope of linear regression with associated standard error, to derive the growth rate in dimensions of days^−1^, as used in Fig. 1E. Raw data provided by Martin Picard.

**Figure S12.**
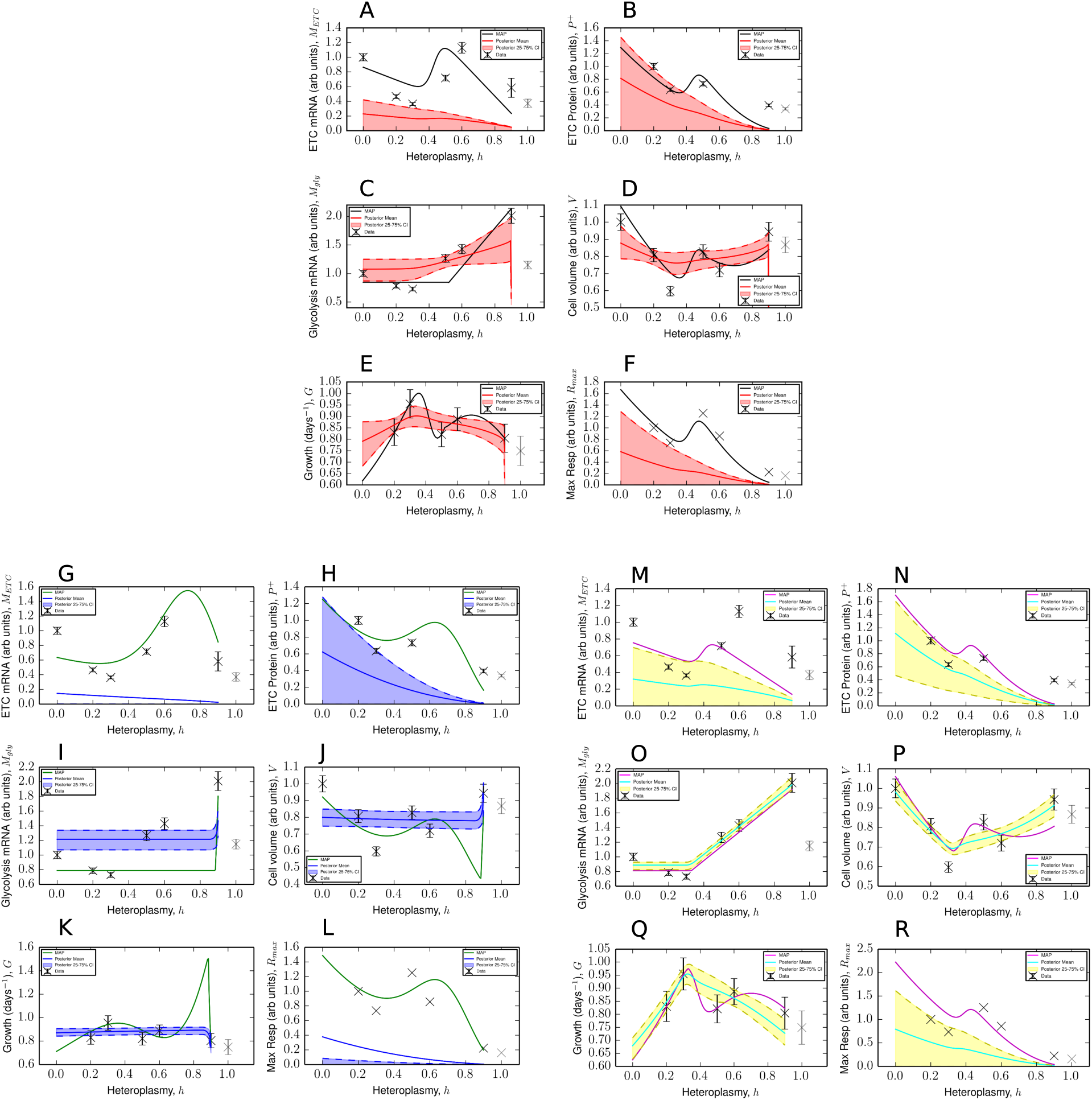
Hyperparameter choice for error model. A-F: Model support for when Ω=0, corresponding to a uniform prior on the variance of each feature. G-L: Sub-samples of posterior where 0.85 ≤ *h*^*\**^ ≤ 0.9. M-R: Sub-samples of posterior where 0.3 ≤ *h*^*\**^ ≤ 0.35. We see a physiologically implausible fit for 0.85 ≤ *h*^*\**^ ≤ 0.9, whereas when 0.3 ≤ *h*^*\**^ ≤ 0.35 model fits were better able to describe the data (for instance, by comparing G to M).

**Figure S13.**
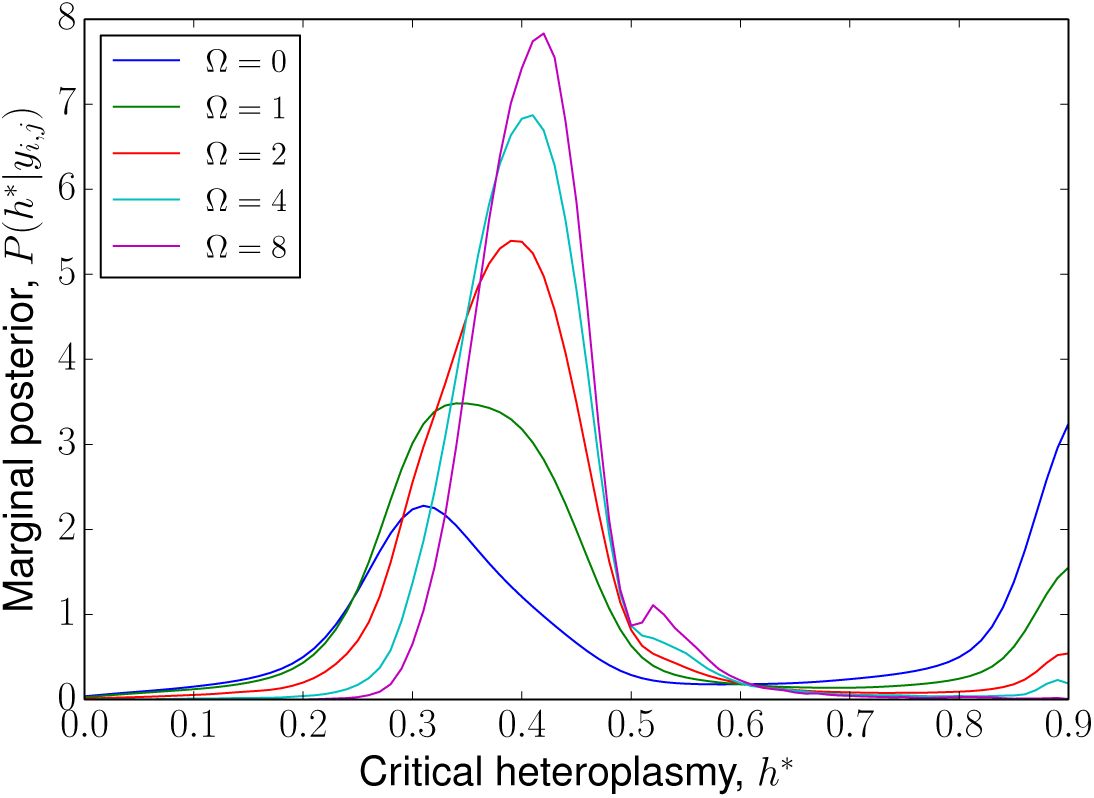
Choice of uncertainty prior affects distribution of critical heteroplasmy and model fit. Larger values of Ω suppresses large values of model uncertainty (*σ*_*i*_, see Eq.(15)), and consequently forces the model fit to more closely match the data. This corresponds to the *h*^*\**^ mode approximately between 0.3-0.4 see Figure S12.

**Figure S14.**
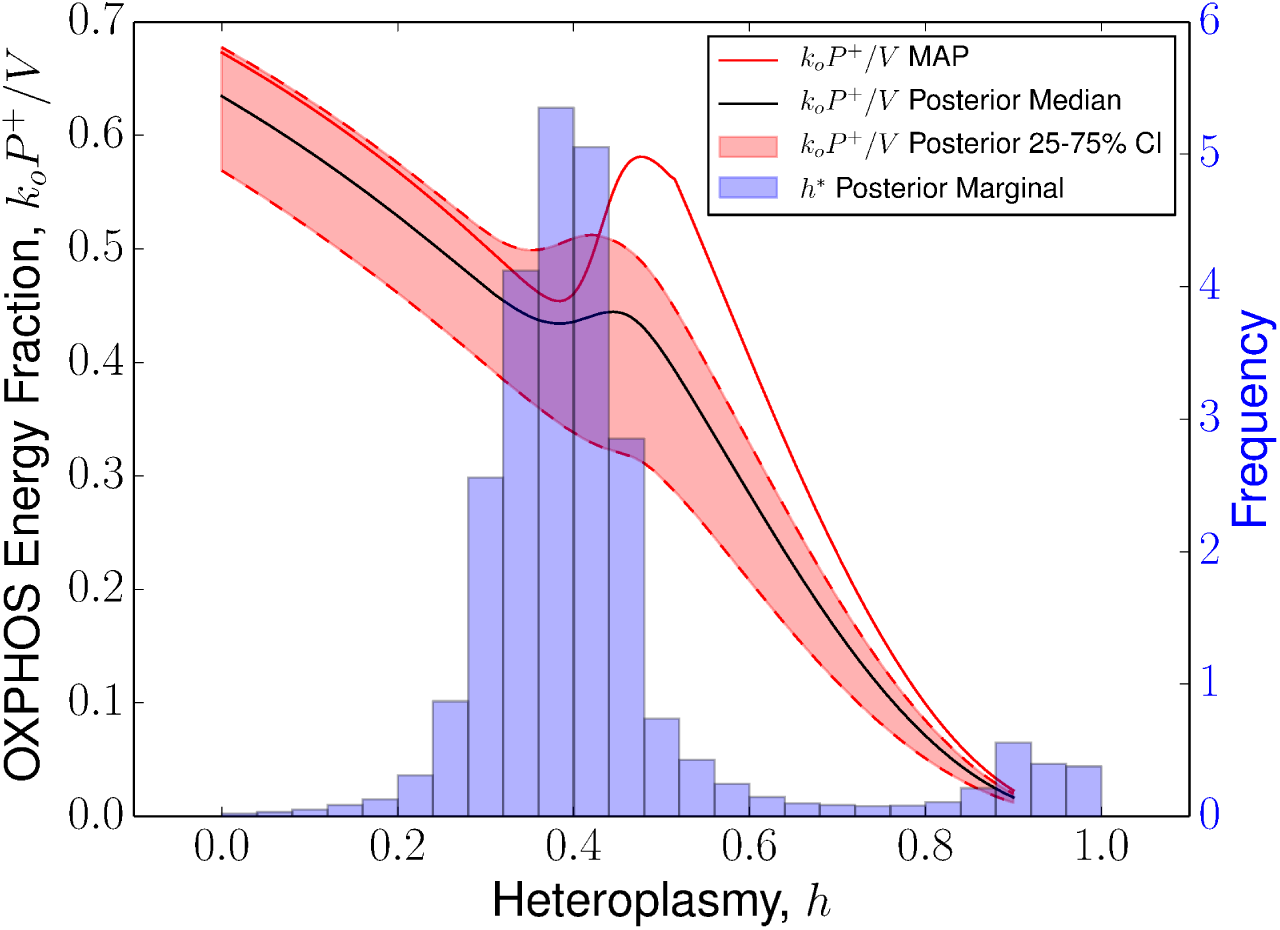
Relative contribution of OXPHOS to total energy supply across heteroplasmy. Posterior statistics for the ratio of OXPHOS energy supply (*k_o_P* ^+^) to total energy supply *k_o_P* ^+^ + *k_g_M*_gly_=*V*. The contribution of ETC power production reduces until the critical heteroplasmy *h*^*\**^, where a compensatory response stabilizes OXPHOS contributions. As 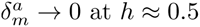 (see Fig. 6), OXPHOS energy contributions continue to diminish.

**Figure S15.**
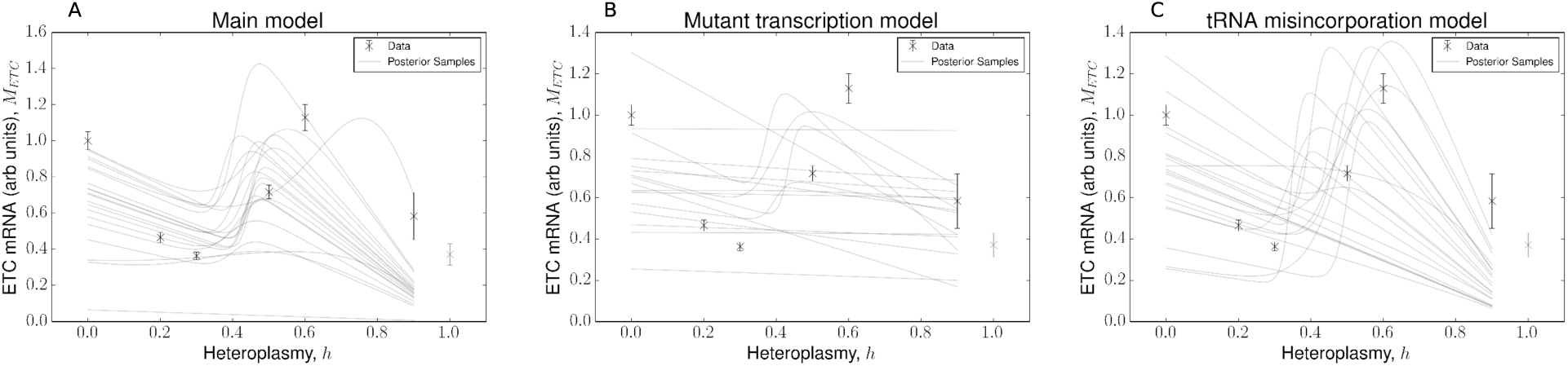
Comparison of ETC mRNA levels for alternative models. Sample of 20 randomly sampled trajectories from the posterior distribution of *M*_*ETC*_ for A. Main model (see Interactions between Bioenergetic Variables can be Cast as a Bottom-up Quantitative Model); B. Mutant transcription model (see Text S4); C. tRNA misincorporation model (see Text S4). In models (B) and (C), we find that linear fits are more frequently selected when compared to our main model (A), despite being more complex in terms of number of parameters.

**Table S1.** Summary of experimental proposals, corresponding to the claims of the model.

| Claim | Experiment |
| --- | --- |
| MtDNA copy number affects volume | Perturb mtDNA copy number (ddC or PGC1-α), measure volume |
| Wild-type mtDNA density affects $h^*$ (bioenergetic) | Increase nDNA content, perform RNA-seq |
| Wild-type mtDNA density affects $h^*$ (mechanical) | Reduce cell volume, perform RNA-seq |
| Mutant mtDNAs have transcription defect | Measure normal/mutant tRNA abundance with qPCR |
| Mitochondrial tRNAs have low diffusivity | Fluorescent labelling of mtDNA and tRNA or mRNA encoding tRNA |
| Cell cycle variation with heteroplasmy | Fucci markers in heteroplasmic cells |
| Mean cell volume affects growth rate | Synchronise, sort by volume, and measure growth rate |
| Maximum respiratory capacity ∝ ETC protein | Measure $R_{max}$ at $h = 0$ and $P^+$ at $h = 0.6$ |

**Table S2.**
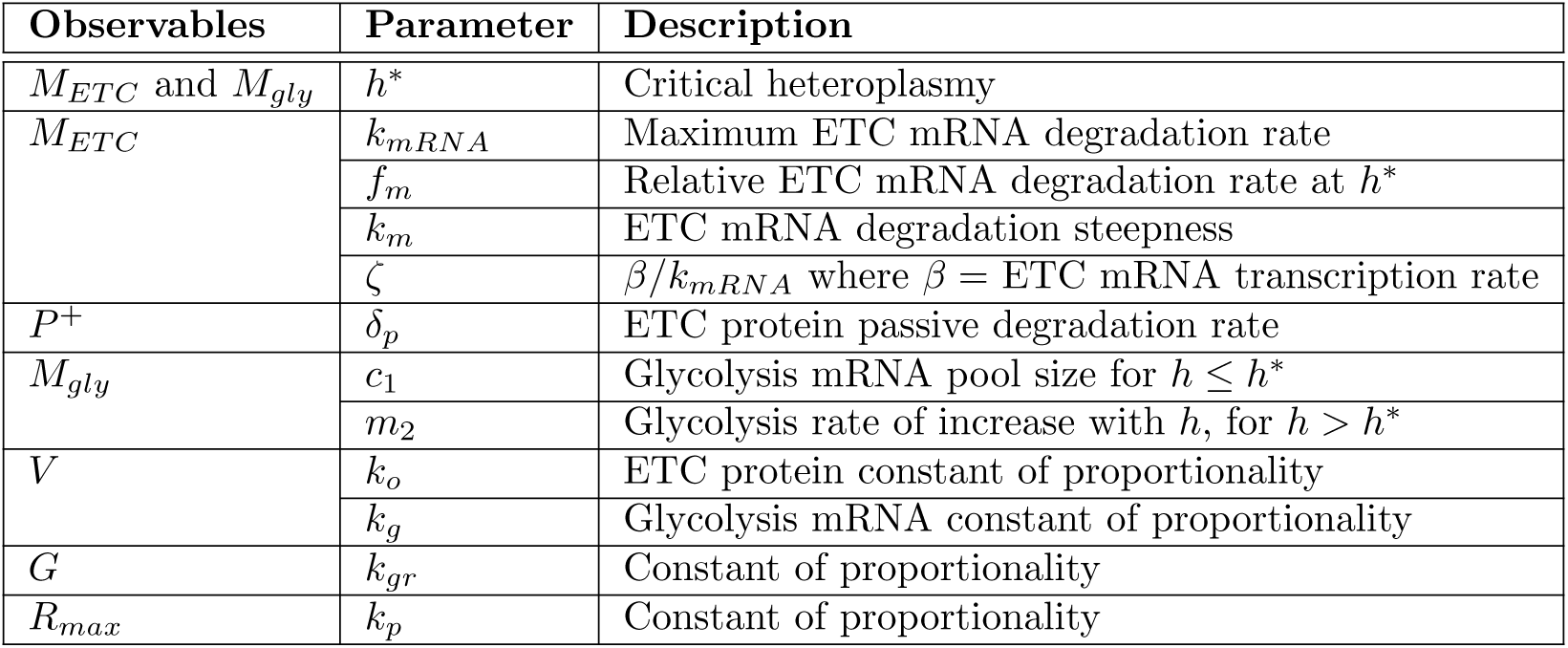
Table of observables, and their corresponding parameters

| Observables | Parameter | Description |
| --- | --- | --- |
| $M_{ETC}$ and $M_{gly}$ | $h^*$ | Critical heteroplasmy |
| $M_{ETC}$ | $k_{mRNA}$ | Maximum ETC mRNA degradation rate |
| | $f_m$ | Relative ETC mRNA degradation rate at $h^*$ |
| | $k_m$ | ETC mRNA degradation steepness |
| | $\zeta$ | $\beta/k_{mRNA}$ where $\beta$ = ETC mRNA transcription rate |
| $P^+$ | $\delta_p$ | ETC protein passive degradation rate |
| $M_{gly}$ | $c_1$ | Glycolysis mRNA pool size for $h \leq h^*$ |
| | $m_2$ | Glycolysis rate of increase with $h$, for $h > h^*$ |
| $V$ | $k_o$ | ETC protein constant of proportionality |
| | $k_g$ | Glycolysis mRNA constant of proportionality |
| $G$ | $k_{gr}$ | Constant of proportionality |
| $R_{max}$ | $k_p$ | Constant of proportionality |

## Acknowledgements

We are grateful to Martin Picard for providing raw data, advice on our model, and experimental suggestions. Till Hoffmann provided technical advice for the Bayesian inference aspects of our work. We would like to thank David Fell, Thomas Ouldridge and Hanne Hoitzing for their useful comments and suggestions.

